# Plant-specific and conserved mechanisms of the polymerase-associated factor 1 complex in replication stress responses

**DOI:** 10.64898/2026.02.09.704960

**Authors:** Cunliang Li, Yuyu Guo, Ziying Wang, Haowei zheng, Jing Deng, Shunping Yan, Lili Wang

**Affiliations:** Hubei Hongshan Laboratory, Huazhong Agricultural University, Wuhan, 430070, China

**Author notes:** Correspondence: Lili Wang. These authors contributed equally to this work (Cunliang Li, Yuyu Guo).

**Keywords:** Genome stability, DNA replication stress, PAF1, WEE1, RFC

## Abstract

DNA replication stress threatens genome stability in eukaryotes. The evolutionarily conserved kinase WEE1 is essential for the activation of the replication stress response. Although the polymerase-associated factor 1 complex (PAF1C) is highly conserved in eukaryotes, its role in the DNA replication stress response remains unclear. Here, we show that Arabidopsis PAF1C is essential for replication stress response. PAF1C-deficient mutants exhibit hypersensitivity to hydroxyurea (HU)-induced replication stress. Mechanistically, we uncover a plant-specific regulatory pathway in which WEE1 interacts with and phosphorylates the PAF1 subunit within its unique N-terminal domain, thereby preventing PAF1 polyubiquitination and subsequent proteasomal degradation to ensure PAF1 accumulation under stress. Genetically, overexpression of a phospho-mimetic PAF1 variant suppresses the HU hypersensitivity of *wee1*, revealing that PAF1 acts downstream of WEE1. However, the WEE1-PAF1 regulatory axis is absent in yeast, indicating its lineage-specific innovation. Further studies reveal that the replication factor C (RFC) complex interacts with and recruits PAF1 to the stalled replication forks. PAF1 then sequentially recruits the E2 ubiquitin-conjugating enzymes UBC1/2 and the E3 ubiquitin ligases HUB1/2 to promote histone H2B monoubiquitination (H2Bub), thereby facilitating replication fork stability. This RFC-dependent recruitment mechanism is conserved in yeast. Collectively, this study suggests that PAF1 regulates replication stress responses by integrating a plant-specific protein stability control mechanism (WEE1-PAF1) with a conserved recruitment mechanism (RFC-PAF1-UBC1/2-HUB1/2), uncovering a novel function of PAF1C and revealing new mechanisms of WEE1 and the RFC complex.

## Introduction

Accurate DNA replication is the foundation of cell proliferation. However, various endogenous and exogenous factors often disrupt this process, causing replication forks to slow, stall, or even collapse. This is known as DNA replication stress ^1^. If replication stress is not resolved properly, it will lead to genome instability ^2^, causing various diseases such as cancers in mammals and developmental defects in plants ^3,4^. To cope with these threats, organisms have evolved sophisticated replication stress response mechanisms, including arresting the cell cycle, stabilizing the replication fork, and stimulating dNTP biosynthesis ^5,6^.

The protein kinase WEE1 is a central and conserved regulator of replication stress response in eukaryotes. In animals, replication stress activates WEE1 by its upstream kinase ataxia telangiectasia and rad3-related (ATR). The activated WEE1 then inhibits cyclin-dependent kinases (CDKs), via phosphorylation, thereby blocking cell cycle progression and ensuring sufficient time for DNA repair ^7–10^. However, this mechanism is not conserved in plants. It has been demonstrated that WEE1 functions independently of its role in phosphorylating CDKA;1, the sole classical CDK that contains the conserved PSTAIRE motif in Arabidopsis thaliana ^11^. We recently found that the loss of the E3 ubiquitin ligase F-box-like 17 (FBL17), the MOS4-associated complex (MAC) subunit pleiotropic regulatory locus 1 (PRL1), or the translational inhibitor general control nonderepressible 20 (GCN20) inhibits the hypersensitivity of the *wee1* mutant to replication stress. FBL17, PRL1, and GCN20 promote cell cycle progression through distinct mechanisms: FBL17 inhibits the CDK inhibitors (CKIs); PRL1 regulates the splicing of cell cycle genes; and GCN20 inhibits the translation of suppressor of gamma response 1 (SOG1), a master transcription factor that activates the expression of CKIs in response to replication stress. Under DNA replication stress, the activated WEE1 phosphorylates FBL17, PRL1, and GCN20, promoting their polyubiquitination and subsequent degradation, which results in an increase in the activity of CKIs, the splicing defects of some cell cycle genes, and the enhanced translation of SOG1, thereby arresting the cell cycle ^12–14^. In addition, the Lieven group reported that mutations in the fasciata1 (FAS1) subunit of the chromatin assembly factor-1 (CAF-1) or in DNA polymerase α suppress the phenotypes of *wee1* ^15,16^. These findings indicate that plant WEE1 functions through multiple mechanisms, unlike its counterparts in other eukaryotes. More plant WEE1 substrates remain to be discovered.

The replication factor C (RFC) complex is a highly conserved heteropentameric ATPase complex in eukaryotes and is required for DNA replication and DNA damage response ^17,18^. The RFC complex is comprised of one large subunit, RFC1, and four small subunits, RFC2/3/4/5 ^19–23^. RFC1 can be substituted by RAD17 to form an alternative form of RFC, the RFC-like complex (RLC) ^24^. During DNA replication and repair, the RFC complex functions as a clamp loader to load the sliding clamp proliferating cell nuclear antigen (PCNA), the key component of the replication machinery, and serves as a processivity factor of DNA polymerases, onto the primer-template junctions in an ATP-dependent manner ^18,25–27^. Recent studies have demonstrated that the depletion of RFC and PCNA results in the de-protection of DNA ends and DNA replication fork collapse ^28^. However, the precise mechanism by which the plant RFC complex regulates the replication stress response remains unclear.

The polymerase-associated factor 1 (PAF1) complex (PAF1C) is also evolutionarily conserved in all eukaryotes. The widely recognized function of PAF1C is to regulate transcription by directly interacting with RNA polymerase II (RNA Pol II) and regulating chromatin modification ^29^. PAF1C can recruit E2 ubiquitin-conjugating enzymes UBC1/2 and E3 ubiquitin ligases HUB1/2 to mediate the monoubiquitination of H2B (H2Bub) that is critical for the dimethylation or trimethylation of H3K4, H3K36, and H3K79 ^30–32^. In Arabidopsis, PAF1C consists of six subunits, PAF1, VIP3, VIP4, VIP5, VIP6, and CDC73 ^33–35^. It was reported that Arabidopsis PAF1C regulates multiple critical biological processes. PAF1 inhibits flowering by promoting the expression of *FLC* through an interaction with RNA Pol II ^36,37^. Recently, we demonstrated that PAF1C facilitates DNA double-strand break (DSB) repair by being recruited to DSB sites through SMC5/6 ^38^, and that PAF1C controls the growth-defense tradeoff by repressing defense gene expression through its interaction with histone deacetylases ^39^. Nevertheless, its precise function in the replication stress response remains unclear.

In this study, we establish PAF1C as a key regulator in the replication stress response. WEE1 phosphorylates PAF1, thereby inhibiting its polyubiquitination and degradation to enhance its protein stability. Furthermore, the RFC-like complex recruits PAF1 to the replication forks, where PAF1 promotes H2Bub by sequentially recruiting UBC1/2 and HUB1/2, thereby stabilizing the replication forks. More importantly, we found that WEE1 regulation of PAF1 is plant-specific, whereas the RFC-like complex recruitment of PAF1 is evolutionarily conserved. Thus, these findings not only reveal the mechanisms by which PAF1 and the RFC complex regulate the replication stress response but also provide a clear example of a core biological function (replication stress response) being accomplished by the same components (WEE1, PAF1, and RFC) but with differentiated regulatory mechanisms evolved across different species.

## Results

### The PAF1 complex is involved in replication stress responses

We previously identified the *ddrm4-1* mutant through a forward genetic screen for DNA Damage Response Mutants (DDRM), and *s*ubsequently confirmed that this mutant exhibits PAF1 functional defects ^38^. In addition to exhibiting hypersensitivity to DSB-inducing agents ^38^, we found that the *paf1-1 (ddrm4-1)* and *paf1-2 (ddrm4-2)* mutants also displayed significant hypersensitivity in root growth upon exposure to hydroxyurea (HU), a replication stress-inducing agent that inhibits ribonucleotide reductase (Fig. 1a and 1b). Furthermore, the root length of the complementation lines (*COM*), in which the genomic sequence of *PAF1* driven by its native promoter (*pPAF1:PAF1*) was transformed into *paf1-1,* was similar to that of Col-0 in the presence of HU (Fig. S1), indicating that PAF1 fully complements the *paf1* mutant under replication stress. These results suggested that PAF1 is involved in the replication stress response.

**Fig. 1.**
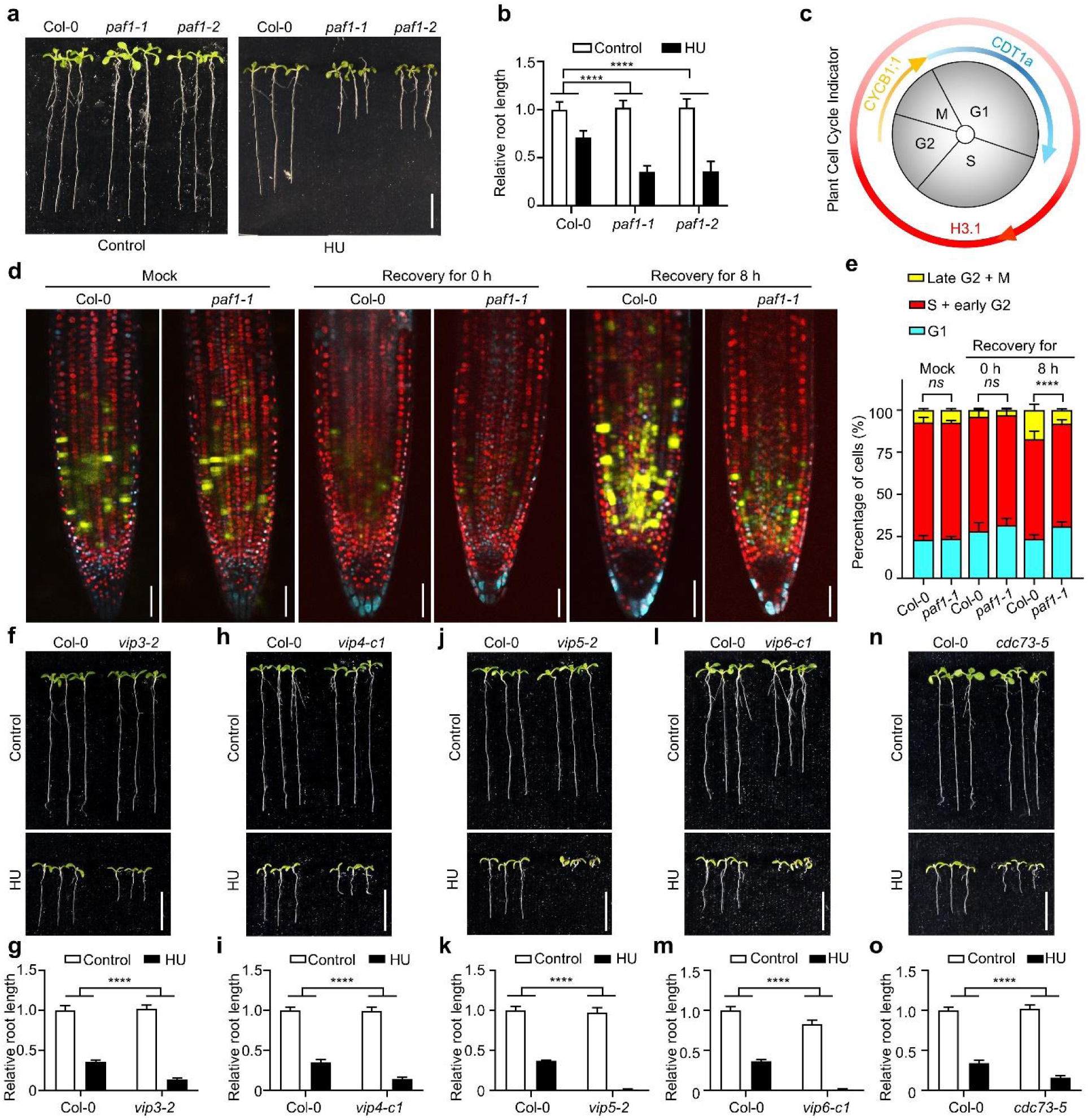
PAF1C is involved in the replication stress response. **a, f, h, j, l, and n,** Pictures of plants treated with HU. Plants were grown vertically on 1/2 MS medium with or without 1 mM HU for 7-8 days. Scale bar = 1 cm. **b, g, i, k, m, and o,** The relative root length of the indicated plants. The data are represented as means ± SD (n = 10 plants) relative to the values obtained under the control conditions. The statistical significance was determined using two-way ANOVA analysis. *\*\*\*\*P* < 0.0001. **c,** Schematic representation of cell cycle phases, color-coded according to the fluorescence expression window of the plant cell cycle indicator (PlaCCI) line. Phase proportions are not to scale. Cyan shows CDT1a-CFP signal, indicating that the cells are in the G1 phase. Red shows H3.1-mCherry signal, indicating that the cells are in the S and early G2 phases. Yellow shows CYCB1;1-YFP signal, indicating that the cells are in the late G2 and M phases. **d,** Representative images of roots of the *PlaCCI/Col-0* and *PlaCCI/paf1-1* plants. Seedlings were treated with or without HU for 4 h and then transferred to 1/2 MS medium for recovery. Roots were imaged by confocal microscopy after 0 and 8 h of recovery. Scale bar = 50 μm. **e,** Quantification of cell percentages at different cell cycle stages. The data are represented as the mean percentage of cells at each cell cycle stage ± SD (n = 4 roots, > 1923 cells total). The statistical significance was determined using two-way ANOVA analysis for the interaction factor (cell cycle stage × genotype). *ns*, not significant; *\*\*\*\*P* < 0.0001.

To investigate the impact of PAF1 deficiency on cell cycle progression under replication stress, we employed the plant cell cycle indicator (PlaCCI) line ^40–42^, which expresses three cell cycle markers (CDT1a–CFP, H3.1–mCherry, and CYCB1;1–YFP) that selectively label cells at different cell cycle stages (Fig. 1c). We crossed the PlaCCI line with *paf1-1* to observe the distribution of cell cycle stages within the root cells. The seedlings were treated with HU and then transferred to 1/2 MS medium for recovery. When treated with HU for 4 h (recovery for 0 h), both Col-0 and *paf1-1* roots showed a significantly reduced proportion of cells in the late G2 and M phases (Fig. 1d and e). After recovery for 8 h, the proportion of cells entering the late G2 and M phases was significantly higher in Col-0 than in *paf1-1* (Fig. 1d and e), indicating that WT roots efficiently resumed the cell cycle, whereas *paf1-1* failed to recover similarly, retaining a higher fraction of cells in the G1 and S phases, likely due to unresolved replication stress. These results suggested that PAF1 is essential for the replication stress response.

PAF1 is the core subunit of PAF1C, which also contains five additional subunits, VIP3, VIP4, VIP5, VIP6, and CDC73 ^43^. To determine whether the hypersensitivity of *paf1* to replication stress is attributed to the specific function of PAF1 or the general function of PAF1C, we examined the phenotypes of the other five subunit mutants generated previously (Li et al 2023). Like the *paf1* mutants, all these mutants exhibited hypersensitivity to HU (Fig. 1f-o), suggesting that the whole PAF1C is involved in replication stress response.

### PAF1 interacts with WEE1

To elucidate how PAF1C regulates the replication stress response, we tested potential interactions between PAF1C and key replication stress regulators (including WEE1, PRL1, FBL17, and SOG1) using yeast two-hybrid (Y2H) assays. Notably, we found that PAF1 interacts with the protein kinase WEE1 (Fig. 2a). Consistent with this finding, the PAF1C subunits PAF1, VIP6, and CDC73 were identified as proximal interactors of WEE1 (Dataset S1 and Fig. S2) through the proximity labeling technique TurboID coupled with mass spectrometry. This analysis was performed in the *pGVG:WEE1-TurboID-HA/wee1* transgenic Arabidopsis, in which dexamethasone (DEX)-inducible expression of WEE1-TurboID-HA partially rescued the HU hypersensitivity of *wee1* (Fig. S2a), confirming functional complementation.

**Fig. 2.**
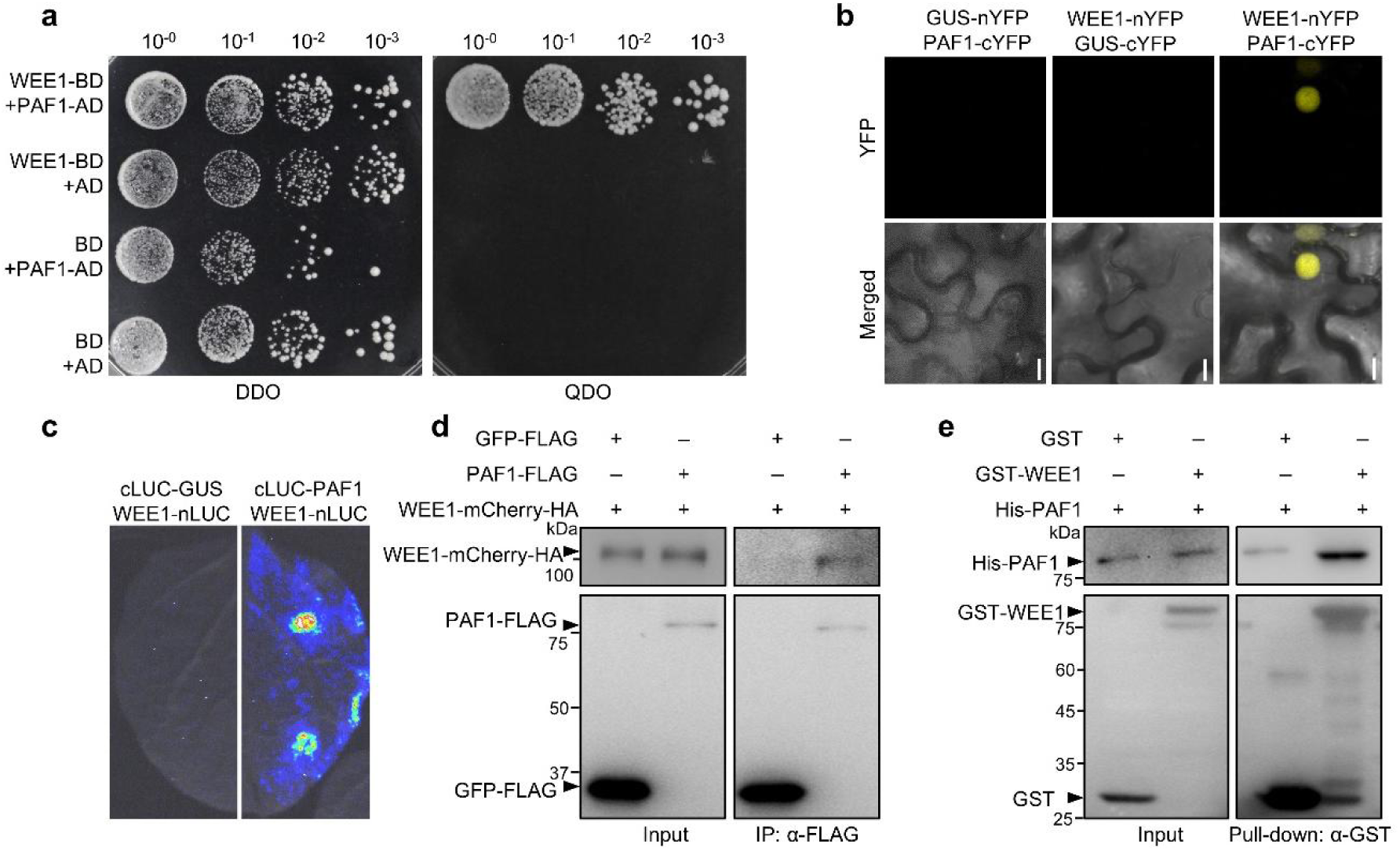
PAF1 interacts with WEE1. **a,** Y2H assays. AD, activation domain. BD, DNA-binding domain. DDO, double dropout (SD/-Trp/-Leu) medium. QDO, quadruple dropout (SD/-Trp/-Leu/-His/-Ade) medium. The yeasts were grown for 3 days. **b,** BiFC assays. The proteins were fused to either the C- or N-terminal half of YFP (cYFP or nYFP) and were transiently expressed in *N. benthamiana.* GUS serves as a negative control. The YFP fluorescence detected by confocal microscopy indicates interaction. Scale bars = 10 μm. **c,** Split luciferase assays. The proteins were fused to either the C- or N-terminal half of luciferase (cLUC or nLUC) and were transiently expressed in *N. benthamiana*. The luminescence detected by a charge-coupled device (CCD) camera indicates interaction. **d,** Co-IP assays. WEE1-mCherry-HA was co-expressed with PAF1-FLAG or GFP-FLAG in Arabidopsis protoplasts. The immunoprecipitation was carried out using anti-FLAG beads. **e,** *In vitro* pull-down assays. The glutathione beads coupled with GST or GST-WEE1 were incubated with His-PAF1, respectively. All experiments were repeated at least three times with similar results.

To further validate the interaction of PAF1 and WEE1 *in planta*, we performed bimolecular fluorescence complementation (BiFC) assays (Fig. 2b), split luciferase assays (Fig. 2c), and co-immunoprecipitation (Co-IP) assays (Fig. 2d). These results revealed that PAF1 specifically interacted with WEE1 *in vivo* and this interaction occurred in the nucleus (Fig. 2b). To determine whether this interaction is direct, we conducted *in vitro* pull-down assays (Fig. 2e). GST-WEE1, but not GST alone, efficiently pulled down more His-PAF1 (Fig. 2e), demonstrating a direct physical interaction. Collectively, these results suggested that PAF1 physically interacts with WEE1 both *in vivo* and *in vitro*.

### WEE1 phosphorylates PAF1 at Y43, Y45, Y56, Y57, and Y63

Given that WEE1 is a functional kinase and interacts with PAF1, we hypothesized that WEE1 could phosphorylate PAF1. To test this, we performed an *in vitro* protein phosphorylation assay ^44^ by co-expressing His-PAF1 with either GST-WEE1 or GST in *E. coli*. The results showed that co-expression with WEE1, but not GST, resulted in the appearance of slower-migrating bands above the main His-PAF1 band (Fig. 3a). To confirm whether these bands represented the phosphorylated forms of PAF1, we analyzed the samples using Phos-tag SDS-PAGE, which retards the migration of phosphorylated proteins by binding to them. The results showed that the multiple shifted bands above His-PAF1 were more clearly separated (Fig. 3a), confirming these bands were phosphorylated forms of PAF1.

**Fig. 3.**
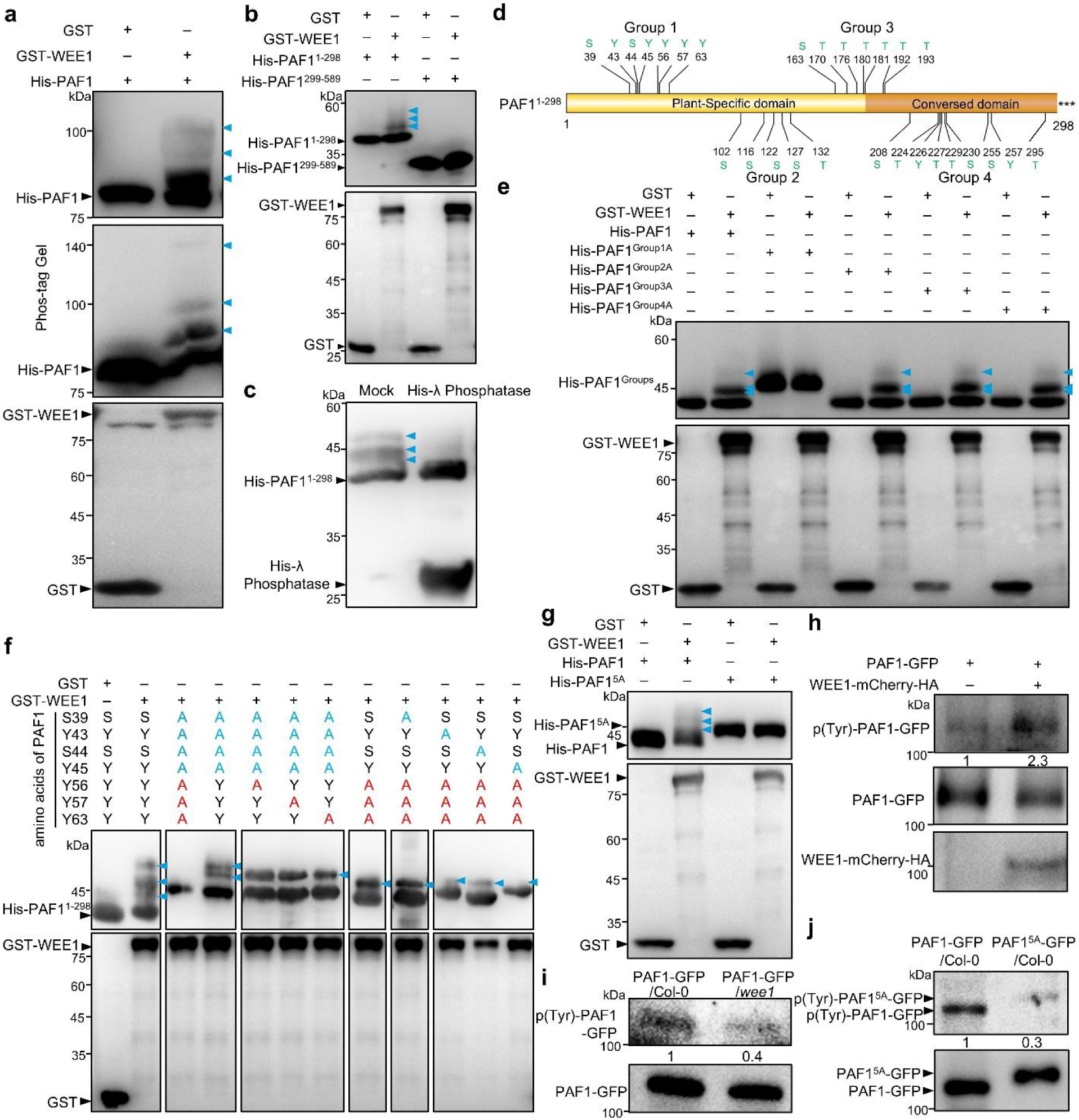
WEE1 phosphorylates PAF1 at Y43, Y45, Y56, Y57, and Y63. **a-c and e-g,** *In vitro* protein phosphorylation assays. GST or GST-WEE1 was co-expressed with different mutant forms of His-PAF1 in the same vector in *E. coli*. The different mutant forms of His-PAF1 were purified using Ni-agarose beads and subjected to immunoblot analysis. The proteins in the middle image of (a) were separated on a Phos-tag gel, while proteins in the other images were separated on SDS-PAGE. The purified His-PAF1^1–298^ protein was treated with or without λ phosphatase for 4 h (c). His-PAF1^Group 1A/2A/3A/4A^ indicates that the amino acids in Group 1/2/3/4 of His-PAF1^1–298^ were mutated to alanine (e). His-PAF1^5A^ represents that the amino acids Y43, Y45, Y56, Y57, and Y63 in His-PAF1 were mutated to alanine (g). **d,** Schematic diagram of PAF1^1–298^. Most of the potential phosphorylation sites (including tyrosine, serine, and threonine residues) are indicated and grouped into 4 groups. **h,** *In vivo* protein phosphorylation assays in protoplasts. PAF1-GFP was co-expressed with or without WEE1-mCherry-HA in *wee1* protoplasts. The protoplasts were treated with 1 mM HU for 4 h before total protein extraction. The PAF1-GFP proteins were immunoprecipitated with anti-GFP beads and analyzed by immunoblot analysis with an anti-Phosphotyrosine antibody. **i and j,** *In vivo* protein phosphorylation assays in transgenic plants. The indicated transgenic plants were treated with 5 mM HU for 4 h. Immunoprecipitation was carried out using anti-GFP beads. The blue arrowheads indicate phosphorylated PAF1. All experiments were repeated at least three times with similar results.

To map the phosphorylation sites of PAF1, we generated two truncated versions of PAF1, PAF1^1–298^ and PAF1^299–589^. The *in vitro* protein phosphorylation assays in *E. coli* showed that the phospho-shifted bands were detected above His-PAF1^1–298^ but not above His-PAF1^299–589^ (Fig. 3b), suggesting that the phosphorylation sites of PAF1 are in its N-terminal 1–298 amino acids. The shifted bands above PAF1^1–298^ were eliminated by the addition of protein phosphatase (Fig. 3c), further confirming that they were phosphorylated bands. To further narrow down the phosphorylation region, we divided most of the potential phosphorylation residues (tyrosine, serine, and threonine) of PAF1 1–298 amino acids into 4 groups (Groups 1–4) based on their spatial distribution (Fig. 3d) and generated His-PAF1^Group 1A^ to His-PAF1^Group 4A^ mutants of His-PAF1^1–298^ in which the potential phosphorylation sites within each group were substituted with alanine (A). These mutated His-PAF1^1–298^ variants were co-expressed with GST-WEE1 or GST in *E. coli,* respectively. Immunoblot analysis showed that only His-PAF1^Group1A^ lost the phosphorylated bands (Fig. 3e), indicating that the phosphorylation sites of PAF1 are located within Group 1. Group 1 contains 7 potential phosphorylation sites: S39, Y43, S44, Y45, Y56, Y57, and Y63 (Fig. 3d). To refine the mapping, we substituted S39, Y43, S44, and Y45 of PAF1^1–298^ with alanine. The *in vitro* phosphorylation assays showed that this variant retained partial phosphorylation (4th lane in Fig. 3f), suggesting that the remaining three amino acids, Y56, Y57, and Y63, may also be phosphorylated. On this basis, further individual mutations at positions Y56, Y57, or Y63 to alanine resulted in a further reduction in phosphorylation levels (5th, 6th, and 7th lanes in Fig. 3f), indicating that all three amino acids can undergo phosphorylation. Furthermore, mutations at positions Y56, Y57, and Y63 in PAF1^1–298^ to alanine retained partial phosphorylation (8th lane in Fig. 3f), suggesting that the remaining four amino acid residues, S39, Y43, S44, and Y45, may be phosphorylated. Similarly, further individual alanine mutations at positions S39, Y43, S44, and Y45 revealed that mutations at Y43 or Y45 (10th and 12th lanes in Fig. 3f), but not S39 or S44 (9th and 11th lanes in Fig. 3f), further reduced phosphorylation levels, indicating that Y43 and Y45 are phosphorylation sites. Collectively, these data identified five phosphorylation sites of PAF1: Y43, Y45, Y56, Y57, and Y63. To validate these findings, we generated His-PAF1^5A^ in which all five phosphorylation sites (Y43, Y45, Y56, Y57, and Y63) of the full-length PAF1 were replaced with alanine. The *in vitro* phosphorylation assays showed that His-PAF1^5A^ no longer exhibited detectable phospho-bands (Fig. 3g), further confirming these five tyrosine residues as the primary phosphorylation targets of WEE1.

To rule out the possibility that mutations of these five tyrosine residues disrupt the PAF1-WEE1 interaction, thereby abolishing PAF1 phosphorylation, we examined the interaction between the phospho-null PAF1^5A^ or the phospho-mimetic PAF1^5D^ (these five tyrosine residues were replaced with aspartate (D)) and WEE1 by *in vitro* pull-down assays. The result showed that both forms interacted robustly with WEE1 (Fig. S3), suggesting that the phosphorylation status of PAF1 does not affect the PAF1-WEE1 interaction.

To further demonstrate that WEE1 phosphorylates PAF1 *in planta*, we performed *in vivo* phosphorylation assays. PAF1-GFP was co-expressed with or without WEE1-mCherry-HA in *wee1* protoplasts, immunoprecipitated using anti-GFP beads, and analyzed by immunoblot analysis with anti-Phosphotyrosine antibody. As shown in Fig. 3h, the phosphorylation level of PAF1-GFP was higher in the presence of WEE1 than in its absence, suggesting that PAF1 is phosphorylated by WEE1. Consistent with this, in stable transgenic plants, the PAF1-GFP phosphorylation level was significantly lower in *wee1* than that in WT (Fig. 3i). To validate the five phosphorylation sites *in vivo*, we generated the *PAF1-GFP/Col-0* and *PAF1^5A^-GFP/Col-0* transgenic plants, in which the cauliflower mosaic virus (CaMV) *35S* promoter drives the expression of the fusion proteins. The *in vivo* phosphorylation assays showed that PAF1^5A^-GFP exhibited negligible phosphorylation bands compared to PAF1-GFP (Fig. 3j), suggesting that WEE1-mediated phosphorylation of PAF1 *in vivo* depends on these five amino acid residues.

### WEE1 inhibits PAF1 polyubiquitination and proteasomal degradation

Our biochemical data demonstrated that WEE1 phosphorylates PAF1, prompting us to investigate whether WEE1 affects PAF1 protein stability, similar to its regulation of the substrates FBL17, PRL1, and GCN20 ^12–14^.

Since WEE1 activity is controlled by replication stress, we first investigated whether replication stress affects PAF1 protein levels. As shown in Fig. S4a and 4b, HU treatment significantly enhanced the fluorescence intensity of PAF1-GFP in transgenic plants. Consistently, neither transiently expressed PAF1-FLAG in Arabidopsis protoplasts nor PAF1-FLAG in transgenic plants showed increased protein levels upon HU treatment (Fig. S4c and d), indicating that HU elevates PAF1 protein levels. To exclude the possibility that the elevated PAF1 protein levels under HU treatment resulted from the increased *PAF1* mRNA expression, potentially caused by HU-mediated modulation of the *35S* promoter activity, we carried out RT-qPCR analysis. The results showed that HU treatment had no significant effect on the mRNA levels *of PAF1 fusions* (Fig. S5a-c). Importantly, we also found that HU treatment did not affect the transcription levels of *PAF1*, as confirmed by RT-qPCR analysis (Fig. S5d) and β-glucuronidase (GUS) staining (Fig. S5e) in the *proPAF1:GUS* transgenic seedlings that the *GUS* gene was driven by the *PAF1* promoter.

Then we hypothesized that replication stress may promote the protein level of PAF1 by inhibiting its degradation. To test this, we carried out semi-*in vitro* protein degradation assays. Recombinant His-PAF1 proteins were incubated with the protein extracts of Col-0 callus treated with or without HU. As shown in Fig. S6a, the degradation of His-PAF1 was slower in the presence of HU than in its absence, indicating that HU inhibits PAF1 degradation. Notably, the degradation of PAF1 could be blocked by the proteasome inhibitor MG132, suggesting that PAF1 is degraded by the ubiquitin-proteasome system. Thus, we investigated whether HU affects the polyubiquitination of PAF1. In the semi*-in vitro* ubiquitination assays, recombinant His-PAF1 proteins coupled with Ni-agarose beads were incubated with the protein extracts of Col-0 callus treated with or without HU. Compared to the control, the polyubiquitination level of His-PAF1 was significantly lower in HU-treated samples (Fig. S6b). To confirm this result *in vivo*, we transiently expressed PAF1-GFP in Arabidopsis protoplasts in the presence or absence of HU. The PAF1-GFP proteins immunoprecipitated using anti-GFP beads were subjected to immunoblot analysis. As shown in Fig. S6c, the polyubiquitination level of PAF1-GFP was significantly reduced in HU-treated samples. Collectively, these results suggested that replication stress inhibits the polyubiquitination and degradation of PAF1.

Next, we investigated whether WEE1 mediates this effect. The semi-*in vitro* ubiquitination assays showed that the PAF1 polyubiquitination level in the *wee1* mutant was higher than that in Col-0 (Fig. 4a). The *in vivo* ubiquitination assays revealed that co-expression of WEE1-mCherry-HA significantly reduced the PAF1 polyubiquitination level compared to PAF1-GFP alone expression in *wee1* protoplasts (Fig. 4b). Thus, these results indicated that WEE1 inhibits PAF1 polyubiquitination. Furthermore, the semi-*in vitro* degradation assays showed that the degradation rate of His-PAF1 in *wee1* was faster than that in Col-0 (Fig. 4c), suggesting that WEE1 inhibits the degradation of PAF1. To further validate this, we carried out co-expression assays in Arabidopsis protoplasts. PAF1-FLAG was co-expressed with different amounts of WEE1-mCherry-HA in *wee1* protoplasts. CFP-HA in the same vectors of PAF1-FLAG was used as a control for transfection efficiency. As shown in Fig. 4d, the PAF1-FLAG protein level was gradually increased when expression of WEE1-mcherry-HA was gradually increased, indicating that WEE1 inhibits PAF1 degradation and thereby promotes its protein levels. Taken together, these results indicated that WEE1 inhibits the polyubiquitination and degradation of PAF1.

**Fig. 4.**
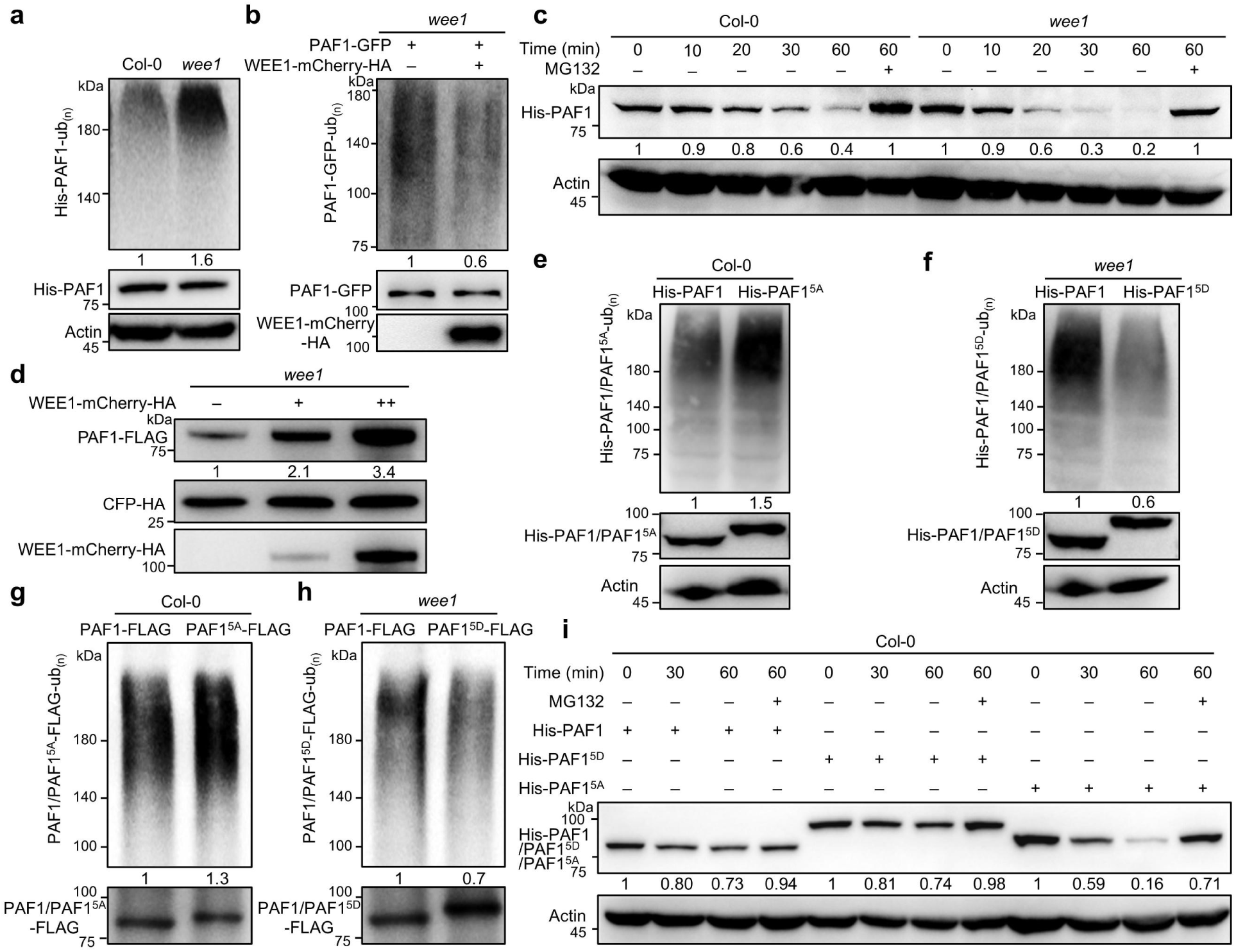
WEE1 inhibits PAF1 ubiquitination and degradation. **a, e, and f,** Semi-*in vitro* ubiquitination assays. The recombinant His-PAF1, His-PAF1^5A^, or His-PAF1^5D^ proteins bound to Ni-agarose beads were incubated with the total protein extracts from HU-treated Col-0 or *wee1* callus in ubiquitination buffer for 4 h. After washing, His-PAF1, His-PAF1^5A^, or His-PAF1^5D^ proteins were subjected to immunoblot analysis using anti-Ubiquitin (Ub) antibody. **b, g, and h,** *In vivo* ubiquitination assays. PAF1-GFP was co-expressed with or without WEE1-mCherry-HA in the *wee1* protoplasts (b). PAF1-FLAG, PAF1^5A^-FLAG, or PAF1^5D^-FLAG was expressed in the Col-0 (g) or *wee1*(h) protoplasts. The protoplasts were treated with 50 μM MG132 and 1 mM HU for 4 h, and then immunoprecipitation was performed using anti-GFP beads or anti-FLAG beads, followed by immunoblot analysis. **c and i,** Semi-*in vitro* protein degradation assays. The recombinant His-PAF1, His-PAF1^5A^ or His-PAF1^5D^ proteins were incubated with total protein extracts of Col-0 or *wee1* callus treated with 5 mM HU. **d,** Co-expression assays in Arabidopsis protoplasts. PAF1-FLAG was co-expressed with or without WEE1-mCherry-HA in the *wee1* protoplasts. CFP-HA in the same vector of PAF1-FLAG was used as the control for transfection efficiency. The total proteins were subjected to immunoblot analysis. All experiments were repeated at least three times with similar results.

To explore the effect of WEE1-mediated phosphorylation on PAF1 ubiquitination and degradation, we compared the ubiquitination and degradation of His-PAF1 and the phospho-null His-PAF1^5^ᴬ or the phospho-mimetic PAF1^5D^. The *semi-in vitro* ubiquitination assays showed that His-PAF1^5A^ exhibited higher polyubiquitination levels than His-PAF1, while His-PAF1^5D^ showed lower levels than His-PAF1 (Fig. 4e and 4f). Consistently, the *in vivo* ubiquitination assays showed similar results (Fig. 4g and h). These results suggested that WEE1-mediated phosphorylation inhibits PAF1 polyubiquitination. Furthermore, the semi-*in vitro* protein degradation assays showed that compared to His-PAF1, His-PAF1^5A^ degraded more rapidly, whereas His-PAF1^5D^ showed no significant difference (Fig. 4i), suggesting that WEE1-mediated phosphorylation inhibits PAF1 degradation. Taken together, these results suggested that WEE1-mediated phosphorylation enhances PAF1 stability by inhibiting PAF1 polyubiquitination and degradation processes.

### PAF1 functions downstream of WEE1

The above biochemical data indicated that PAF1 functions downstream of WEE1. To validate their genetic relationship, we generated the *wee1 paf1-1* double mutant by a genetic cross. As shown in Fig. 5a and b, both *paf1-1* and *wee1* displayed hypersensitivity to HU compared to Col-0. Notably, *wee1* was more sensitive to HU than *paf1-1,* indicating that WEE1 may regulate multiple downstream components, with PAF1 being one of the key components. The *wee1 paf1-1* double mutant exhibited significantly stronger sensitivity to HU than either single mutant. This additive genetic interaction indicated that PAF1’s function is only partially dependent on WEE1. We propose that in *wee1*, disruption of all WEE1 downstream pathways, including PAF1, results in a more severe phenotype. In contrast, *paf1-1* contains only the *paf1* mutation, other downstream pathways regulated by WEE1 can partially maintain function, leading to a milder phenotype. In the *wee1 paf1* double mutant, not only are all WEE1-dependent pathways disrupted, but even the function of PAF1 that might exist independently of WEE1 is lost, resulting in the most severe phenotype. Thus, these data indicated that WEE1 acts upstream of PAF1 and exerts its effects by regulating multiple downstream pathways, with PAF1 serving as one of the key regulatory nodes.

**Fig. 5.**
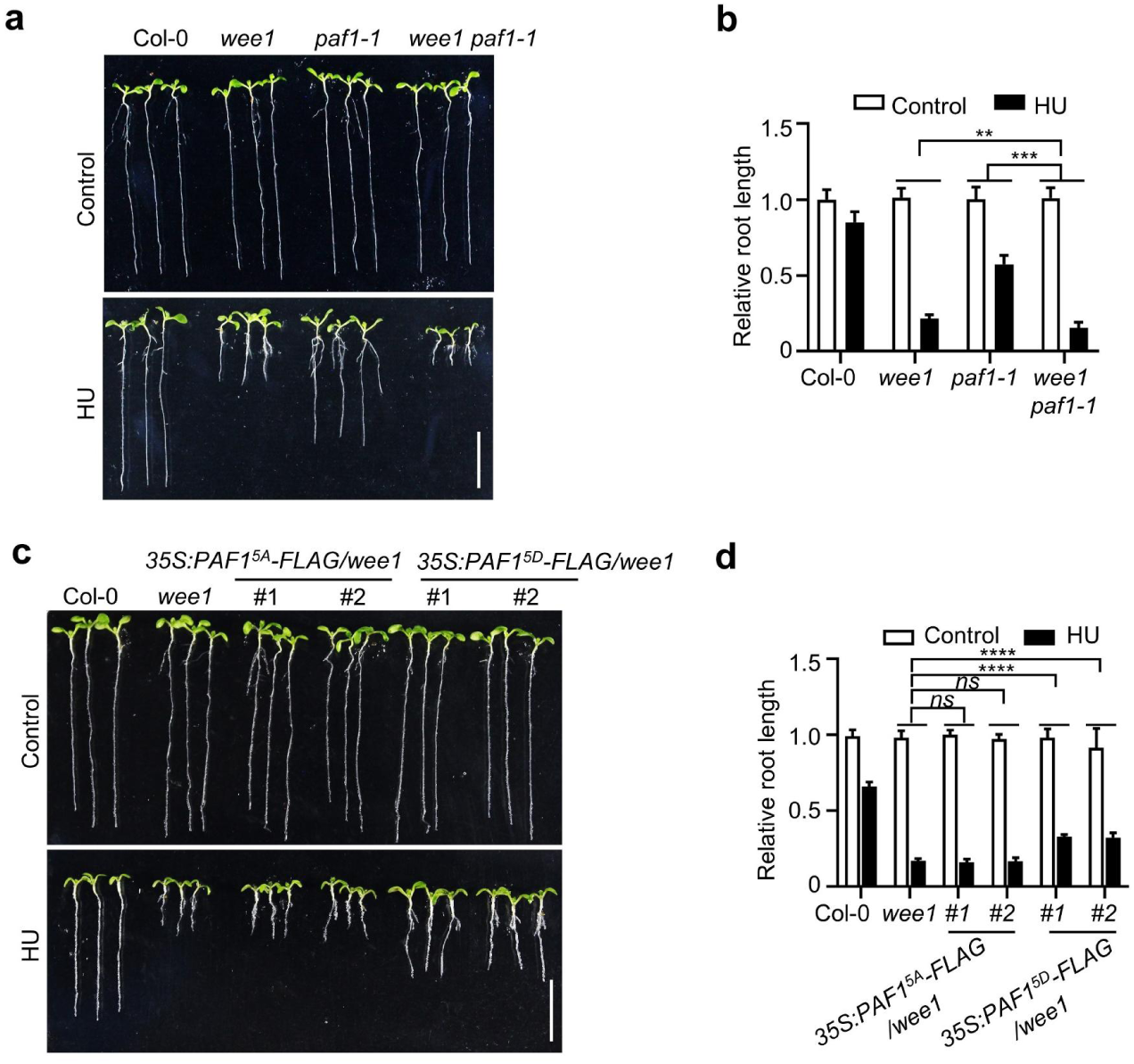
PAF1 functions downstream of WEE1. **a and c,** Pictures of plants treated with HU. Plants were grown vertically on 1/2 MS medium with or without 1 mM HU for 7-8 days. Scale bar = 1 cm. **b and d,** The relative root length of the indicated plants. The data are represented as means ± SD (n = 10 plants) relative to the values obtained under the control conditions. The statistical significance was determined using two-way ANOVA analysis. *ns*, not significant; *\*\*P* < 0.01; *\*\*\*P* < 0.001; *\*\*\*\*P* < 0.0001.

We next sought to investigate the relationship between WEE1 and PAF1 phosphorylation genetically. PAF1^5A^-FLAG and PAF1^5D^-FLAG were overexpressed in the *wee1* mutant, respectively. Strikingly, overexpression of PAF1^5D^-FLAG partially rescued the HU hypersensitivity of *wee1*, whereas PAF1^5A^-FLAG did not (Fig. 5c and d). Moreover, overexpression of PAF1-GFP also failed to rescue the HU hypersensitivity of *wee1* (Fig. S7). Thus, these results further confirmed that PAF1 functions downstream of WEE1, and WEE1 exerts its function by phosphorylating PAF1.

### Evolutionary divergence of the WEE1-PAF1 regulatory module

Given that both WEE1 and PAF1 are conserved across eukaryotes, we sought to investigate whether the regulatory relationship between them is conserved. We first analyzed PAF1 protein sequences. Multiple sequence alignment of PAF1 from 25 representative eukaryotes revealed that PAF1 consists of two regions: an N-terminal region that is specific to plants (Plant-Specific domain, amino acids 1–194) and a C-terminal domain that is widely conserved (Conserved domain, amino acids 195–589) (Dataset S2, Fig. S8a). Phylogenetic analysis confirmed that the N-terminal domain is a plant-specific evolutionary acquisition (Fig. S8a). Importantly, all five phosphorylation sites of PAF1 by WEE1 are located within its N-terminal Plant-Specific domain (Fig. S8a), implying that only plants might use WEE1 to control PAF1. We tested this idea. First, we verified whether the Plant-Specific domain mediates WEE1-PAF1 interaction. Indeed, the split luciferase (Fig. S8b), BiFC (Fig. S8c), and *in vitro* pull-down (Fig. S8d) assays all showed that the Plant-Specific domain, but not the Conserved domain, mediated its interaction with WEE1. We then checked the WEE1-PAF1 interaction in yeast. The *in vitro* pull-down assays showed that *S. cerevisiae* Paf1 (ScPaf1) could not interact with ScWee1 (Fig. S8e). Consistent with this, the *in vitro* phosphorylation assays suggested that ScWee1 could not phosphorylate ScPaf1 (Fig. S8f). These results indicated that the regulatory mechanism by which WEE1 binds to and phosphorylates PAF1 is a plant-specific invention and is not conserved in yeast.

Interestingly, the *scpaf1* mutants generated by replacing *ScPaf1* with orotidine-5’-phosphate decarboxylase (*ODCase*/*URA*) gene in yeast (Fig. S9a and b) exhibited hypersensitivity to HU (Fig. S9c), indicating that ScPaf1 plays a role in the yeast replication stress response. Together, these results suggested that the role of PAF1 in replication stress tolerance is ancient and conserved. The fact that PAF1’s role is old, but the way plants control it is new, led us to explore other ancient ways PAF1 might work.

### Conserved interaction between PAF1 and the RFC-like complex

To elucidate how PAF1 functions during replication stress, we examined a previously published co-purification dataset of PAF1 ^43^ and searched for replication-associated interacting proteins. Several subunits of the RFC complex attracted our attention (Fig. S10) because the RFC complex plays an important role in maintaining genome stability. The RFC complex consists of one large subunit, RFC1, and four small subunits, RFC2/3/4/5 ^19^. RAD17 could replace RFC1 to form the RFC-like (RAD17/RFC2-5) complex ^24^. To validate the interaction between PAF1 and the RFC/RFC-like complex, we performed Y2H assays, which revealed that PAF1 could interact with RFC4, RFC5, and RAD17, respectively (Fig. S11a and 6a). These interactions were further substantiated *in planta* by BiFC, split luciferase, and Co-IP assays (Fig. 6b-d and S11b-g). To test whether their interactions are direct or indirect, we carried out *in vitro* pull-down assays using purified recombinant proteins, which showed that their interactions are direct (Fig. 6e and Fig. S11h-i). Therefore, PAF1 directly interacts with RFC4, RFC5, and RAD17 both *in vitro* and *in vivo*. We next mapped the interaction domain on PAF1. Through Y2H, BiFC, split luciferase, and *in vitro* pull-down assays, we found that RFC4 specifically interacts with the conserved domain of PAF1 (PAF1^195–589^) (Fig. S12a-d). This finding prompted us to test whether the PAF1-RFC interaction is evolutionarily conserved. Indeed, the *in vitro* pull-down assays showed that yeast ScPaf1 could directly interact with ScRfc4 (Fig. S8e and S12e). Together, these data established a direct, conserved interaction between PAF1 and the RFC-like complex, suggesting an ancient functional partnership.

**Fig. 6.**
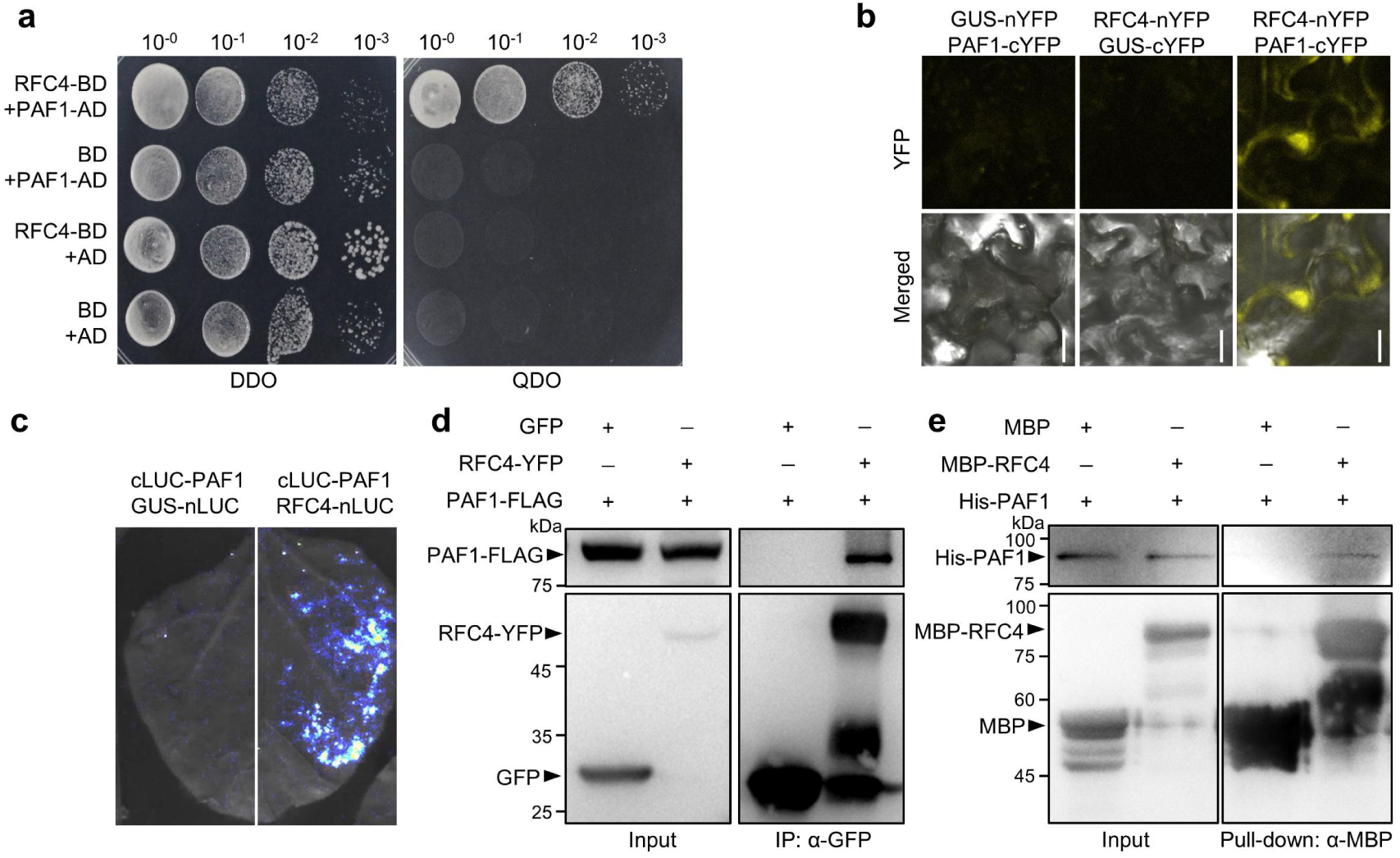
PAF1 interacts with RFC4. **a,** Y2H assays. AD, activation domain. BD, DNA-binding domain. DDO, double dropout (SD/-Trp/-Leu) medium. QDO, quadruple dropout (SD/-Trp/-Leu/-His/-Ade) medium. The yeasts were grown for 3 days. **b,** BiFC assays. The proteins were fused to either the C- or N-terminal half of YFP (cYFP or nYFP) and were transiently expressed in *N. benthamiana*. GUS serves as a negative control. The YFP fluorescence detected by confocal microscopy indicates interaction. Scale bars = 10 μm. **c,** Split luciferase assays. The proteins were fused to either the C- or N-terminal half of luciferase (cLUC or nLUC) and were transiently expressed in *N. benthamiana*. The luminescence detected by a charge-coupled device (CCD) camera indicates interaction. **d,** Co-IP assays. PAF1-FLAG was co-expressed with GFP or RFC4-YFP in Arabidopsis protoplasts. The immunoprecipitation was carried out using anti-GFP beads. **e,** *In vitro* pull-down assays. The MBP-RFC4 or MBP coupled with the Dextrin beads were incubated with His-PAF1. All experiments were repeated at least three times with similar results.

### The RFC-like complex recruits PAF1 to stalled replication forks

Next, we want to know how the PAF1-RFC interaction regulates the replication stress response. We first examined whether replication stress affects their interaction. To this end, we performed HU treatment in the Y2H assays, using the previously reported NPR1-PAF1 interaction as a control ^39^. HU inhibits yeast growth. However, yeasts expressing RFC4 and PAF1 grew better than those expressing NPR1 and PAF1 (Fig. 7a), which is further supported by their growth curves (Fig. 7b). These results suggested that HU promotes the interaction between RFC4 and PAF1. Both BiFC and Co-IP assays *in planta* further confirmed that HU promotes the interaction of PAF1 and RFC4 (Fig. 7c-e).

**Fig. 7.**
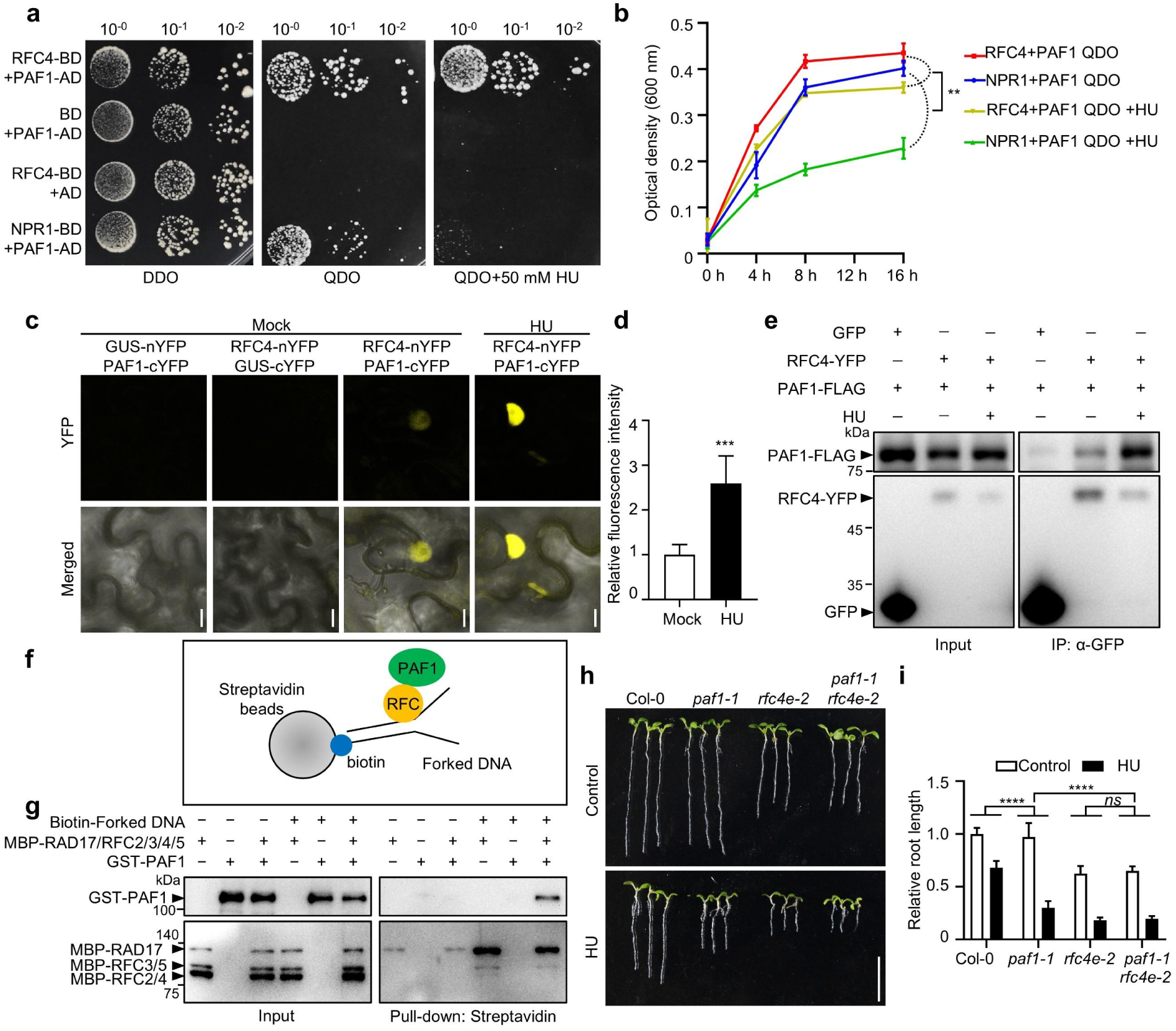
The RFC-like complex recruits PAF1 to the replication fork. **a,** Y2H assays. AD, activation domain. BD, DNA-binding domain. DDO, double dropout (SD/-Trp/-Leu) medium. QDO, quadruple dropout (SD/-Trp/-Leu/-His/-Ade) medium. 50 mM HU was added in the QDO medium. The yeasts were grown for 4 days. **b,** The growth curve of yeast in the liquid QDO medium with or without 50 mM HU. Yeasts co-expressed RFC4 and PAF1, or NPR1 and PAF1. The data are represented as means OD_600nm_ ± SD (n = 3). The statistical significance of the data at 16 h was determined using two-way ANOVA analysis. *\*\*P* < 0.01. **c,** BiFC assays. The proteins were fused to either the C- or N-terminal half of YFP (cYFP or nYFP) and were transiently expressed in *N. benthamiana.* After leaves were treated with or without 5mM HU for 2h, the YFP fluorescence was detected using confocal microscopy. Scale bars = 10 μm. **d,** Statistical analysis of YFP fluorescence intensity in c. The data are represented as means ± SD (n = 6 cells) relative to the values of YFP under the control conditions. The statistical significance was determined using Student’s *t*-test. ***, *P* < 0.001. **e,** Co-IP assays. PAF1-FLAG was co-expressed with GFP or RFC4-YFP in protoplasts with or without HU treatment. The immunoprecipitation was carried out using anti-GFP beads. **f,** Schematic diagram of the forked DNA-binding assays. **g,** Forked DNA-binding assays. Biotin-labeled forked DNA was incubated with GST-PAF1, MBP-RAD17, or/and MBP-RFC2/3/4/5. The forked DNA-binding proteins were pulled down by Streptavidin magnetic beads and analyzed by immunoblot analysis. **h,** Pictures of plants treated with HU. Plants were grown vertically on 1/2 MS medium with or without 1 mM HU for 7-8 days. Scale bar = 1 cm. **i,** The relative root length of the indicated plants. The data are represented as means ± SD (n = 10 plants) relative to the values obtained under the control conditions. The statistical significance was determined using two-way ANOVA analysis. *ns*, not significant; *\*\*\*\*P* < 0.0001. All experiments were repeated at least three times with similar results.

We hypothesized that the RFC-like complex might recruit PAF1 to the stalled replication forks when the replication stress occurs. To test this, we performed forked DNA-binding assays (Fig. 7f) as previously reported ^45^ using purified recombinant GST-PAF1, MBP-RAD17, MBP-RFC2/3/4/5 proteins (Fig. S13), and synthetic biotin-forked DNA. After incubating purified recombinant proteins with biotin-labeled forked DNA, the forked DNA was enriched using streptavidin magnetic beads. Immunoblot analysis of forked DNA-binding proteins showed that the RFC-like complex bound robustly to forked DNA (Fig. 7g). Strikingly, GST-PAF1 was efficiently co-enriched with the DNA forks only in the presence of the RFC-like complex (Fig. 7g), suggesting that the RFC-like complex physically recruits PAF1 to the replication forks.

To test the relationship between PAF1 and the RFC-like complex genetically, we generated the *paf1-1 rfc4e-2* double mutant by a genetic cross. Compared to Col-0, both *paf1-1* and *rfc4e-2* were hypersensitive to HU (Fig. 7h and i). Notably, the HU sensitivity of *paf1-1 rfc4e-2* double mutant was similar to that of *rfc4e-2* (Fig. 7h and i), suggesting that PAF1 and the RFC-like complex function in the same pathway.

### HUB1/2 and UBC1/2 act with PAF1 in the replication stress response

Next, we sought to investigate the functional role of PAF1 after its recruitment to stalled replication forks. Our previous study has shown that PAF1 recruits the E2 ubiquitin-conjugating enzymes UBC1/2 and the E3 ubiquitin ligases HUB1/2 to DSB sites, thereby promoting local histone H2Bub and facilitating DSB repair ^38^. Extensive studies have demonstrated that H2Bub is also critical for maintaining genome integrity during replication stress ^46–48^. It was reported that ScBre1, the homolog of HUB1/2 in yeast, could bind to the replication fork and stimulate local H2Bub to promote DNA repair ^49^. Thus, we hypothesized that PAF1 might recruit UBC1/2 and HUB1/2 to participate in the replication stress response.

We examined the HU phenotypes of the *hub1-5*, *hub2-2,* and *ubc1 ubc2-c1* mutants that were functionally deficient in HUB1, HUB2, and UBC1/2, respectively ^38^. The results showed that all these mutants exhibited clear hypersensitivity to HU (Fig. 8a and b), suggesting that HUB1/2 and UBC1/2 are involved in the replication stress response. To test the relationship between PAF1 and HUB1 genetically, we examined the HU sensitivity of the *paf1-1 hub1-5* double mutant generated previously ^38^. The result showed that *paf1-1 hub1-5* showed HU sensitivity comparable to that of every single mutant (Fig. 8c and d), suggesting that they function in the same pathway.

**Fig. 8.**
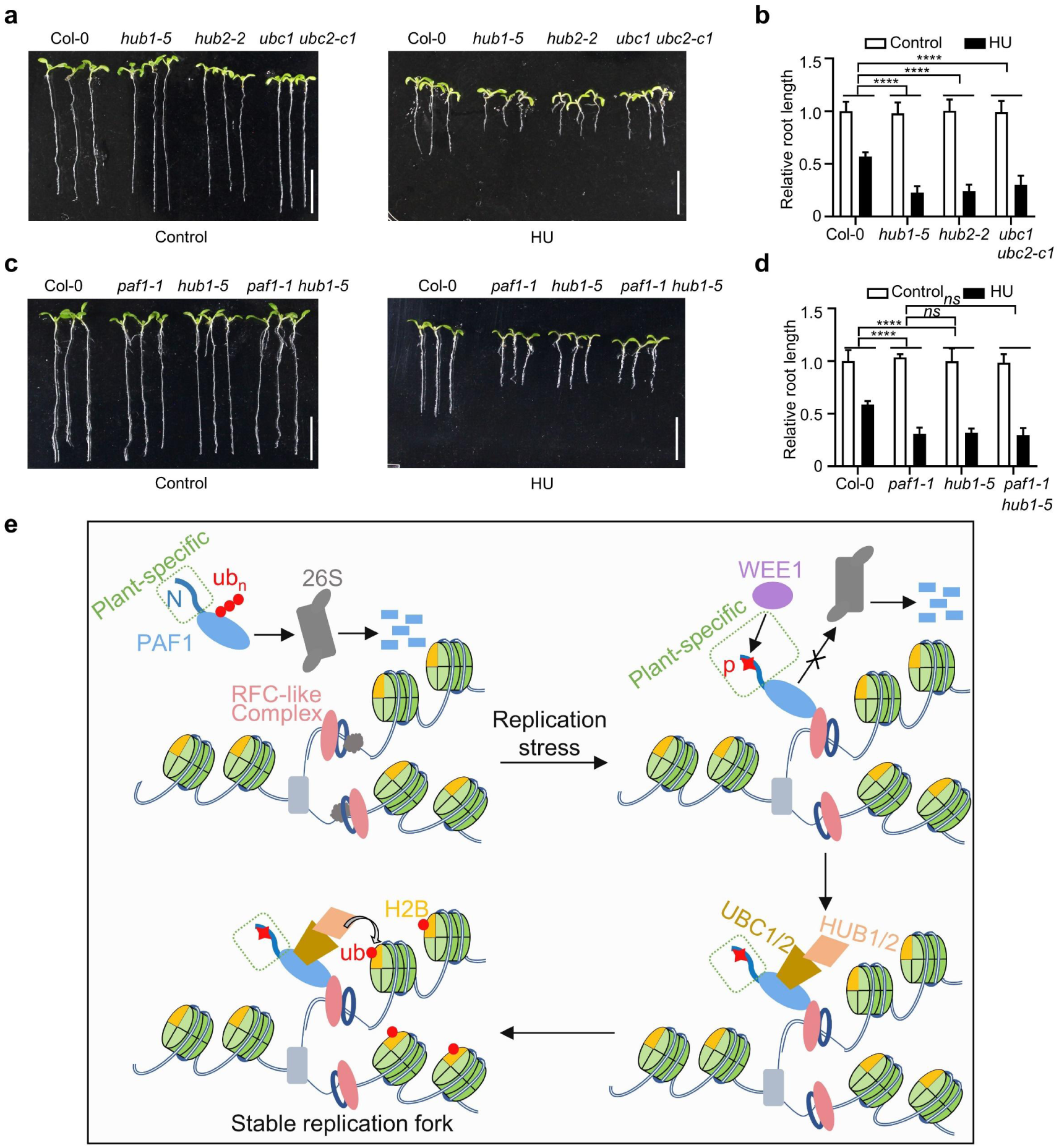
UBC1/2 and HUB1/2 participate in the replication stress response. **a and c,** Pictures of plants treated with HU. Plants were grown vertically on 1/2 MS medium with or without 1 mM HU for 7-8 days. Scale bar = 1 cm. **b and d,** The relative root length of the indicated plants. The data are represented as means ± SD (n = 10 plants) relative to the values obtained under the control conditions. The statistical significance was determined using two-way ANOVA analysis. *ns*, not significant; *\*\*\*\*P* < 0.0001. **e,** A proposed working model to illustrate how PAF1 regulates replication stress responses. “P”, phosphorylation; “ub”, monoubiquitination; ub_n_, polyubiquitination.

## Discussion

Based on the above data, we proposed a working model to illustrate how PAF1 is involved in the replication stress response (Fig. 8e). When replication stress occurs, the protein kinase WEE1 is activated. WEE1 then interacts with the N-terminal Plant-Specific domain of PAF1 and phosphorylates five key tyrosine residues (Y43, Y45, Y56, Y57, and Y63) within this domain. This phosphorylation inhibits PAF1 polyubiquitination and proteasomal degradation, ensuring its rapid accumulation under stress. Concurrently, the evolutionarily conserved RFC-like (RAD17/RFC2-5) complex directly interacts with the conserved domain of PAF1 and efficiently recruits PAF1 to the stalled replication forks. At the fork, PAF1 may further recruit UBC1/2 and HUB1/2 to promote local H2Bub, thereby stabilizing replication forks and maintaining genome stability. Yeast ScWee1 cannot interact with ScPaf1, implying that the WEE1-PAF1 regulatory module is plant-specific and is not conserved in eukaryotes. In contrast, ScRfc4 interacts with ScPaf1, suggesting that the RFC-PAF1 regulatory module is conserved in eukaryotes. Thus, PAF1 function is controlled by a dual regulatory architecture: a plant-specific regulatory pathway that controls PAF1 protein abundance, and an evolutionarily conserved pathway across eukaryotes that directs it to the replication forks.

Notably, although WEE1, PAF1, and the RFC-like complex are broadly conserved in sequence and function across eukaryotes, their regulatory relationships have diverged during evolution. The WEE1-PAF1 stability regulation module is plant-specific, whereas the RFC-PAF1 recruitment mechanism is conserved in eukaryotes. This clearly indicates that during evolution, the core functional capacity of a protein (e.g., PAF1 promotes fork stability) is preserved, but the upstream control layers regulating its activity, abundance, or localization can frequently undergo significant lineage-specific variation. We speculate that the emergence of this plant-specific regulatory module may be linked to the unique selective pressures in sessile plants. For instance, when combating pathogen infection, plants have evolved the immune regulator NPR1, which activates defense responses by directly ubiquitinating PAF1 and promoting its degradation ^39^. Conversely, to tackle replication stress, plants evolved this unique PAF1 stabilization mechanism mediated by WEE1.

In animals and yeast, WEE1 primarily induces cell cycle arrest by regulating CDK activity in response to replication stress. However, plant WEE1 appears to function through a broader set of substrates, including FBL17, PRL1, GCN20, and now PAF1, so that it cannot only coordinate cell cycle arrest, but also coordinate dNTP synthesis, transcriptional reprogramming, and replication fork stabilization. Notably, the regulatory relationships between WEE1 and these substrate proteins are not all conserved across eukaryotes. This further supports the evolutionary principle of “conservation” and “innovation” coexisting. This expansion of WEE1 signaling highlights how the conserved “sentinel” role of a core kinase can be executed through distinct “molecular language” (i.e., different substrates) in different lineages.

This study offers new insights into the roles of WEE1, PAF1, and the RFC complex in replication stress response, but also raises several important questions for future research. First, the boundaries of evolutionary conservation need further clarification. While we have validated the WEE1-PAF1 and RFC-PAF1 interactions in yeast. Future validation in a broader eukaryotic lineage (particularly mammalian cells) is necessary. This will allow us to more precisely map the emergence and divergence of these regulatory modules on the evolutionary tree, thereby deepening our understanding of their evolutionary drivers. Second, the precise mechanism by which PAF1 acts at replication forks remains to be elucidated. Although ScBre1 (the yeast homolog of HUB1/2) has been reported to promote H2Bub at replication forks, and our genetic evidence supports that PAF1 functions in the same pathway as UBC1/2-HUB1/2, direct biochemical and cellular evidence is still lacking. It is unclear whether PAF1 directly recruits these ubiquitination enzymes to the replication fork and promotes local H2Bub in plants. Third, the crosstalk between different stress response pathways requires an integrated study. Our previous work revealed that NPR1 promotes PAF1 degradation for immunity, while this study shows that WEE1 stabilizes PAF1 for genome maintenance. This raises an attractive hypothesis: NPR1 and WEE1 may constitute a molecular switch in plants, balancing resource allocation between immune response and genome maintenance by oppositely regulating PAF1 stability. Exploring whether and how NPR1 participates in the replication stress response will be a crucial next step in understanding how plants prioritize and coordinate responses to different survival threats.

## Materials and Methods

### Plant materials and growth conditions

All Arabidopsis mutants used in this study are in the *Columbia-0* (Col-0) ecotype background. The *vip3-2* (SALK_139885), *cdc73-1* (SALK_150644), *hub1-5* (SALK_044415), and *hub2-2* (SALK_071289) mutants were obtained from Arashare (https://www.arashare.cn). The *paf1-1, ubc1 ubc2-c1, vip5-2* (SALK_062223), *vip4-c1*, *vip6-c1* and *rfc4e-2* mutants were described previously ^38,50,51^. The *paf1-2* (SALK_046605) and *wee1* (SALK_147968C) mutants were obtained from the Arabidopsis Biological Resource Center (ABRC). All the transgenic plants were generated using the floral-dip method ^52^. Seeds were surface-sterilized with 2% Plant Preservative Mixture (PPM^TM^, Plant Cell Technology), stratified at 4°C in the dark for 2 days, and then sown on half-strength Murashige and Skoog (1/2 MS) medium containing 1% (w/v) sucrose and 0.35% (w/v) phytagel (#CP8581Z, Coolaber). Plants were grown under long-day conditions (16 h of light and 8 h of dark) in a growth chamber at 22°C. The calluses were induced on MS medium containing 0.5mg/l 2,4-D and 0.05 mg/l kinetin.

### Vector Construction

All plasmid constructs were generated using the One Step Cloning Kit (#C117, Vazyme Biotech Co.,Ltd). The primers used for cloning are listed in Dataset S3. Coding sequences (CDSs) were amplified and then inserted into the indicated vectors as follows. For Y2H assays, the CDSs of *RFC2/3/4/5*, *RAD17,* and *WEE1* were cloned into *Eco*RI/*Bam*HI-digested *pGBKT7* vector, and the CDSs of *PAF1* was cloned into *Eco*RI/*Bam*HI-digested *pGADT7* vector. For BiFC assays, the CDSs of *RFC4/5*, *RAD17,* and *WEE1* were cloned into *Bam*HI/*Sal*I-digested *pSPYNE-35S* vector, and the CDSs of PAF1 were cloned into *Bam*HI/*Sal*I-digested *pSPYCE-35S* vector. For split luciferase assays, the CDSs of *RFC4/5, RAD17,* and *WEE1* were cloned into the *pJW771* vector digested with *Kpn*I and *Sal*I. The CDSs of PAF1 were cloned into *Kpn*I/*Sal*I-digested *pJW772* vector. For Co-IP assays, the CDSs of *RFC4/5* and *RAD17* fused with a YFP tag were cloned into *Nco*I/*Xba*I-digested *pFGC5941* vector. The CDS of *PAF1* was cloned into *Bam*HI/*Sal*I-digested *pCambia2306* vector. For protein expression in *Escherichia coli*, the CDSs of *RFC4/5, RAD17, ScRfc4*, and *ScWee1* were cloned into *Eco*RI/*Sal*I-digested *pMAL-C2X* vector. The CDSs of *ScPaf1* and *WEE1* were cloned into *Bam*HI/*Not*I-digested *pGEX-6P-1*. The CDSs of PAF1 were cloned into *Bam*HI/*Nde*I -digested *pET28a* vector. For the *in vitro* protein phosphorylation assay. The His-PAF1 and GST-WEE1 were cloned into *pQLink* plasmids. To produce a co-expression plasmid from two pQLink plasmids, one plasmid was digested with *Pac*I at 37°C while the other was cleaved with *Swa*I at 25°C. The digested DNA was treated with T4 DNA polymerase and dCTP or dGTP. Products of the enzyme reaction were mixed and heated to 65°C and cooled to room temperature for annealing. The product was transformed into *E. coli*. Transformants were tested for expected inserts by colony PCR. For GUS staining, the 2-kb promoter fragment upstream of the *PAF1* start codon was cloned into *Bam*HI/*Kpn*I-digested *pCambia2300* vector. For protein expression assays, the CDS of *PAF1* fused with *FLAG* was cloned into *Nco*I/*Pac*I-digested *pFGC5941-CFP-HA* vector.

### RNA extraction and RT-qPCR

Total RNA was extracted using TRIzon Reagent (#CW0580S, CWBIO). The cDNA was synthesized using the First-Strand cDNA Synthesis Kit (#BOLG1025S, Baiuoleji (Hubei) Biotechnology). The Quantitative PCR (qPCR) was performed with Taq SYBR Green qPCR Premix (#BOLG2016M, Baiuoleji (Hubei) Biotechnology) on a Fluorescent Quantitative PCR Detection System (#Q2000B, LongGene). *UBQ5* was used as the internal control. Relative expression levels were calculated using the 2^-ΔΔCT^ method.

### Confocal microscopic observation

For the fluorescence observation of the PlaCCI plants, the seedlings grown vertically on 1/2 MS medium were transferred to 1/2 MS medium containing 1 mM HU for 4 h and then transferred to 1/2 MS medium for 0 or 8 h. For the BiFC assays, the leaves of *N. benthamiana* were treated with 5 mM HU for 2 h. For the fluorescent proteins in transgenic Arabidopsis, the seedlings grown vertically on 1/2 MS medium were transferred to 1/2 MS medium containing 1 mM HU for 4 h. The fluorescence was captured using confocal laser scanning microscopy (TCS SP8, Leica). To avoid fluorescence interference, the sequential scan mode was used.

### Protein interaction analysis

The Y2H, BiFC, split luciferase, Co-IP, and *in vitro* pull-down assays were performed as described previously ^39^. For Y2H assays, the corresponding constructs were co-transformed into the yeast strain AH109. Transformants were selected on appropriate dropout media. For BiFC assays, the fusion proteins were co-expressed in *N. benthamiana*. The YFP fluorescence was observed 48 h post-infiltration using a confocal laser scanning microscope (TCS SP8, Leica). For split luciferase assays, the fusion proteins were co-expressed in *N. benthamiana*. The luminescence signals were captured using the Lumazone Imaging System equipped with a 2048B charge-coupled device camera (Roper). For Co-IP assays, the fusion proteins were transiently co-expressed in Arabidopsis protoplasts. Total protein extracts were incubated with anti-GFP (#L-1016, Biolinkedin) or anti-FLAG beads (#016-101-003, AlpVHHs) at 4°C for 2 h for immunoprecipitation. Beads were washed extensively, and bound proteins were eluted and analyzed by immunoblot analysis. For *in vitro* pull-down assays, all recombinant proteins were expressed in *E. coli* BL21 (DE3). The MBP-tagged proteins coupled to Dextrin beads (#SA026005, Smart-Lifesciences) and GST-tagged proteins coupled to glutathione beads (#SA010005, Smart-Lifesciences) were used to pull down GST-tagged or His-tagged proteins.

### *In vitro* protein phosphorylation assay

The phosphorylation assays using a bacterial reconstituted system were performed as described previously with some modifications ^44^. The *pQLink* co-expression plasmids containing the indicated His-PAF1 variants and GST-WEE1 or GST were transformed into *E. coli* BL21 (DE3). The His-PAF1 variants were purified with Ni-agarose beads (#N30210 Beijing LABLEAD Inc.) and then boiled in the phosphorylation loading buffer [50 mM Tris-HCl (pH 6.8), 1% SDS, 0.1% Bromophenol Blue, 10% glycerine], followed by immunoblot analysis using an anti-His antibody. At the same time, the total bacterial lysates were also boiled in 1× SDS loading buffer, followed by immunoblot analysis using an anti-GST antibody. For phosphatase treatment, the His-PAF1^1–298^ bound to the Ni-agarose beads were incubated with or without His-λ Phosphatase in the 1× λ phosphatase buffer (#P2316S, Beyotime) containing 1 mM MnCl_2_ at 30 °C for 4 h and then boiled in 1× SDS loading buffer, followed by immunoblot analysis using an anti-His antibody.

For the Phos-tag technology, the samples were resolved on 6% SDS-PAGE gels containing 20 μM Phos-tag (#AAL-107, Wako) and 40 μM MnCl_2_ to separate phosphorylated from non-phosphorylated proteins. After electrophoresis, proteins were transferred to membranes and subjected to immunoblot analysis with an anti-His antibody.

### *In vivo* protein phosphorylation assay

Protoplasts were treated with or without 1 mM HU, and transgenic plants were treated with 5 mM HU; all treatments were performed for 4 h. The total proteins were extracted with radioimmunoprecipitation assay (RIPA) lysis buffer [50 mM Tris-HCl (pH 7.5), 150 mM NaCl, 1% Triton X-100, 1% sodium deoxycholate, 0.1% SDS, 1× protease inhibitor, 2 mM phenylmethylsulfonyl fluoride (PMSF), 2 mM dithiothreitol (DTT), and 50 μM MG132], and then incubated with anti-GFP beads at 4°C for 2 h. The beads were washed three times with washing buffer [50 mM Tris-HCl (pH 7.4), 1 M NaCl, 1% Triton X-100, 1% sodium deoxycholate, 0.1% SDS, and 50 μM MG132] and boiled in 1× SDS loading buffer, followed by immunoblot analysis using anti-Phosphotyrosine (#ab10321, Abcam), and anti-GFP (#11814460001, Roche) antibodies.

### Semi*-in vitro* protein degradation assay

The recombinant His-PAF1, His-PAF1^5A^, and His-PAF1^5D^ proteins were incubated at 22°C with the total protein extracts in a protein degradation buffer [25 mM Tris-HCl (pH 7.5), 10 mM NaCl, 10 mM MgCl_2_, 1 mM PMSF, 1 mM DTT, 2 mM adenosine 5′-triphosphate (ATP)] for the indicated times. The total protein extracts were prepared from Col-0 or *wee1* callus treated with or without 5 mM HU for 4 h. Reactions were stopped by adding SDS loading buffer, and proteins were analyzed by immunoblot analysis using anti-His (#2366, Cell Signaling Technology) or anti-actin antibodies.

### Semi*-in vitro* ubiquitination assay

The His-PAF1, His-PAF1^5A^, or His-PAF1^5D^ proteins coupled with Ni-agarose beads were incubated with total protein extracts in ubiquitination buffer [25 mM Tris-HCl (pH 7.5), 10 mM NaCl, 10 mM MgCl2, 2 mM PMSF, 2 mM DTT, 4 mM ATP, 50 μM MG132, and 1× protease inhibitor cocktail] at 22°C for 4 h. The total protein extracts were sampled from Col-0 or *wee1* callus treated with or without 5 mM HU for 4 h. The beads were washed three times with washing buffer and boiled in 1× SDS loading buffer, followed by immunoblot analysis using an anti-ubiquitin antibody. The input samples were subjected to immunoblot analysis using anti-actin or anti-His antibodies.

### *In vivo* ubiquitination assay

Protoplasts were treated with 50 μM MG132 and/or 1 mM HU, and transgenic plants with 5 mM HU; all treatments were performed for 4 h. The total proteins were extracted with RIPA lysis buffer. Lysates were incubated with anti-GFP or anti-FLAG beads at 4°C for 2 h for immunoprecipitation. The beads were washed three times with washing buffer and then boiled in 1× SDS loading buffer, followed by immunoblot analysis using anti-ubiquitin (#3936, Cell Signaling Technology), anti-GFP or anti-FLAG (#11814460001, Roche) antibodies.

### Forked DNA-binding assay

Forked DNA-binding assay was performed as described previously ^45^. All recombinant proteins were expressed in *E. coli* BL21 and purified with Ni-agarose or Dextrin beads. To obtain Biotin-Forked DNA, the 10 μM primers (biotin-GGGCGGCGGGTTTTTTTTTTTTTTTTTTTT and CCCGCCGCCCTTTTTTTTTTTTTTTTTTTT) were mixed and heated to 95°C for 20 min and cooled to room temperature for annealing. The Forked DNA was attached to magnetic beads (#HY-K0208, MCE) via a biotin-streptavidin linkage. The beads were incubated in a stepwise fashion with the MBP-RFC2/3/4/5, MBP-RAD17, and His-PAF1 proteins in the reaction buffer [25 mM Tris-HCl (pH 7.5), 10 mM NaCl, 10 mM MgCl_2_, 1 mM PMSF, 1 mM DTT, 2 mM adenosine 5′-triphosphate (ATP)]. The beads were washed three times with washing buffer [10 mM Tris-HCl (pH 7.5), 1 mM EDTA, 1 M NaCl, 0.1% Tween-20] and boiled in 1× SDS loading buffer. Bound proteins were detected by immunoblot analysis using anti-MBP or anti-His antibodies.

## Acknowledgments

We are grateful to Dr. Yong Ding for providing the *vip5-2* mutant, Dr. Shitou Xia for providing the *rfc4e-2* mutant, Dr. Jiafu Long for providing yeast strains SEY6210, Dr. Yanli Xiang and Dr.Xuelei Lai for critical reading of the manuscript.

## Funding

This work was supported by the National Natural Science Foundation of China (32570353, 32270306, and 32070312) and Huazhong Agricultural University Scientific & Technological Self-innovation Foundation (2662025SKPY005).

## Author contributions

L.W., C.L., and Y.G. designed the experiments; C.L., Y.G., Z.W., J.D, and H.Z. carried out the experiments; Y.G., C.L., S.Y., and L.W. wrote the manuscript; C.L. visualized the data. All authors discussed the results and commented on the manuscript.

## Competing interests

The authors declare no competing interests.

## Notes

### Competing Interest Statement

The authors have declared no competing interest.

