## Supplementary material for "Plant-specific and conserved mechanisms of the polymerase-associated factor 1 complex in replication stress responses": Supplemantal information: ! WEE1-PAF1-supplemental-V11.pdf

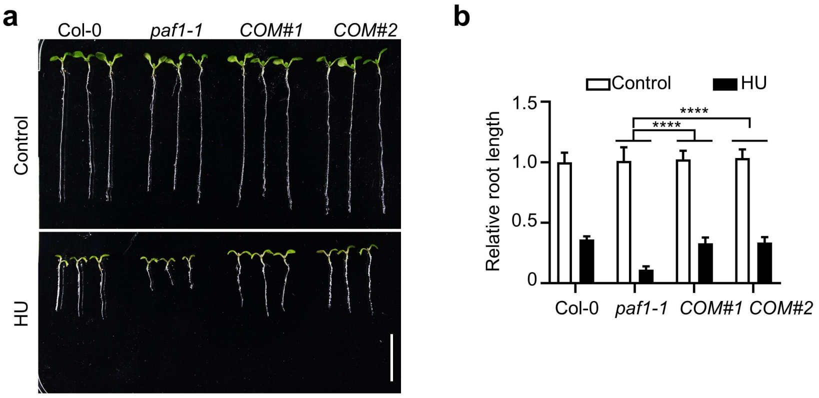


**Fig. S1 PAF1 fully complements the hypersensitivity of *paf1* mutant to replication stress.**

**a,** Pictures of plants treated with HU. Plants were grown vertically on 1/2 MS medium with or without 1 mM HU for 7 days. *COM*, the complementation lines expressing PAF1 driven by its native promoter in *paf1-1*. Scale bar = 1 cm. **b,** The relative root length of the indicated plants. The data are represented as means ± SD (n = 10 plants) relative to the values obtained under the control conditions. The statistical significance was determined using two-way ANOVA analysis. ****, $P < 0.0001$.

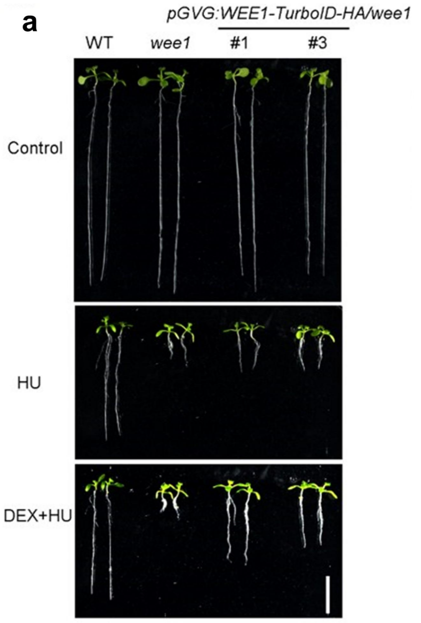


b

TurboID-Mass Spectrometry of WEE1 (Partial data )

| Protein | MS/MS count | Score |
|---|---|---|
| PAF1 | 5 | 18.874 |
| VIP6 | 4 | 26.290 |
| CDC73 | 2 | 13.455 |

**Fig. S2 WEE1 co-purified with PAF1.**

**a,** Pictures of *pGVG:WEE1-TurboID-HA/wee1* transgenic Arabidopsis under different growth conditions. Plants were grown vertically on 1/2 MS medium with or without 1 mM HU and 10 μM dexamethasone (DEX) for 9 days. Scale bar = 1 cm. **b,** Partial TurboID-Mass Spectrometry (MS) results of WEE1.

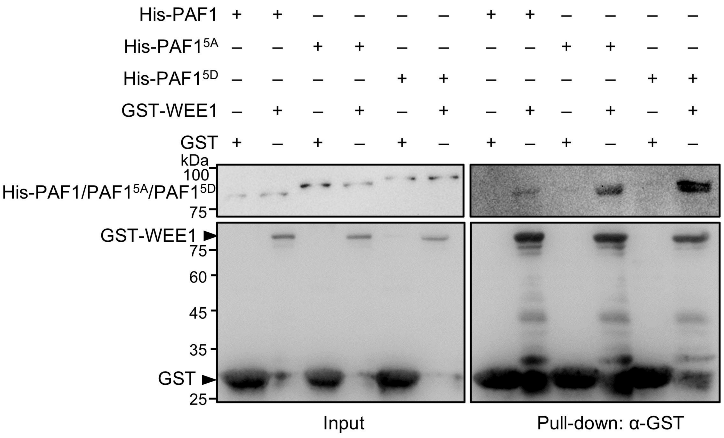


**Fig. S3 PAF1$^{5A}$ and PAF1$^{5D}$ still interact with WEE1 in the *in vitro* pull-down assays.**

The glutathione beads coupled with GST or GST-WEE1 were incubated with His-PAF1, His-PAF1$^{5A}$, or His-PAF1$^{5D}$, respectively.

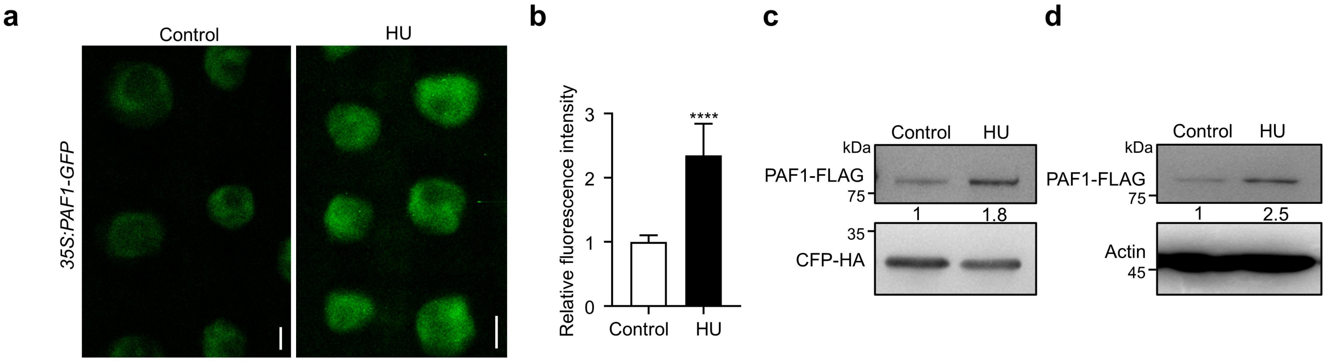


**Fig. S4 HU promotes PAF1 accumulation.**

**a,** Representative images of PAF1-GFP in the *35S:PAF1-GFP/*Col-0 transgenic Arabidopsis treated with or without 1 mM HU for 4 h. Scale bar = 5 μm. **b,** The relative fluorescence intensity of PAF1-GFP in (a). The data are represented as means ± SD relative to the values of PAF1-GFP under the control conditions. The statistical significance was determined using a Student's *t*-test. ****, $P < 0.0001$. **c and d,** The protein levels of PAF1-FLAG in protoplasts (c) and *35S:PAF1-FLAG/Col-0* transgenic Arabidopsis (d) treated with or without 1 mM HU for 4 h. CFP-HA in the same vector of PAF1-FLAG was used as the control for transfection efficiency. The total proteins were subjected to immunoblot analysis.

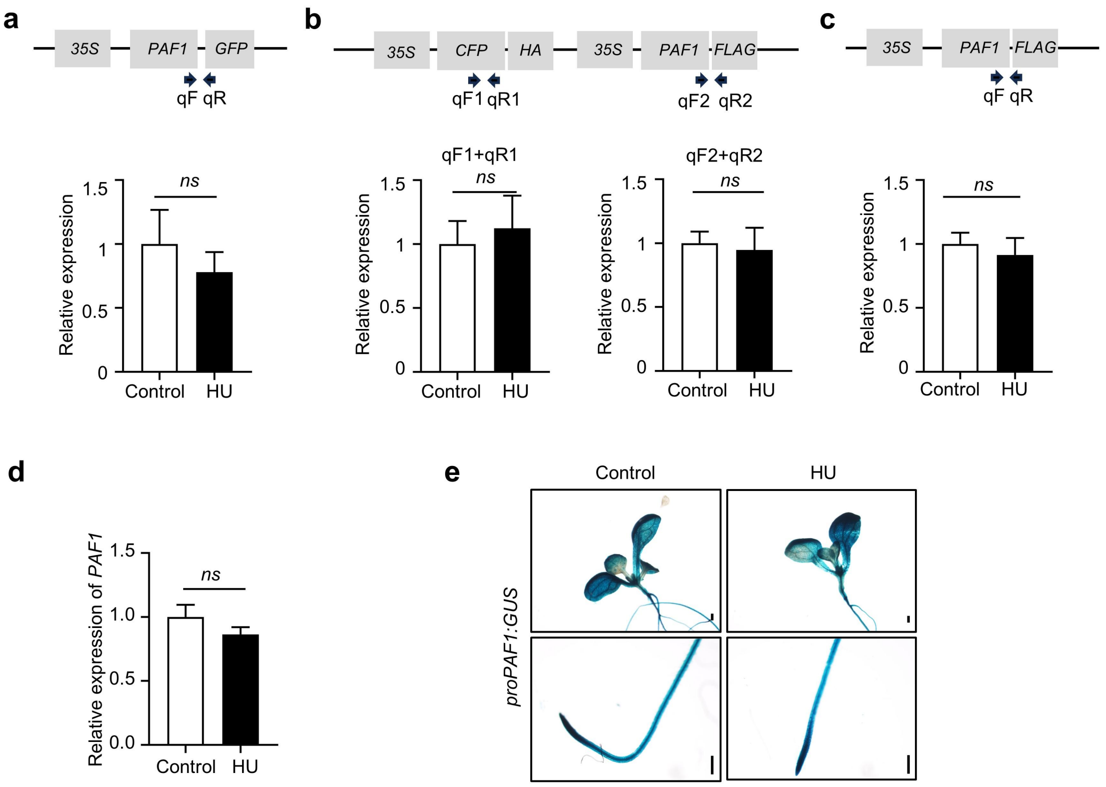


**Fig. S5 HU does not affect *PAF1* transcription.**

**a-c,** The relative expression of *PAF1* fusion genes determined by RT-qPCR analysis. The primers used for RT-qPCR are indicated by black arrows. a shows the relative expression of *PAF1-GFP* in Fig. S4a, b shows the relative expression of *PAF1-FLAG* and *CFP-HA* in the protoplasts in Fig. S4c, and c shows the relative expression of *PAF1-FLAG* in the *35S:PAF1-FLAG* transgenic Arabidopsis in Fig. S4d. *UBQ5* was used as the reference gene. The data are represented as means ± SD (n = 3). The statistical significance was determined using a Student's *t*-test. *ns*, not significant. **d,** Relative expression of *PAF1* determined by RT-qPCR analysis. The Col-0 seedlings were treated with or without 5 mM HU for 4h. *UBQ5* was used as the reference gene. The data are represented as means ± SD (*n* = 3). The statistical significance was determined using a Student's *t*-test. *ns*, not significant. **e,** GUS staining results of the *proPAF1:GUS* seedlings treated with or without 5 mM HU for 4 h. Scale bar = 2 mm.

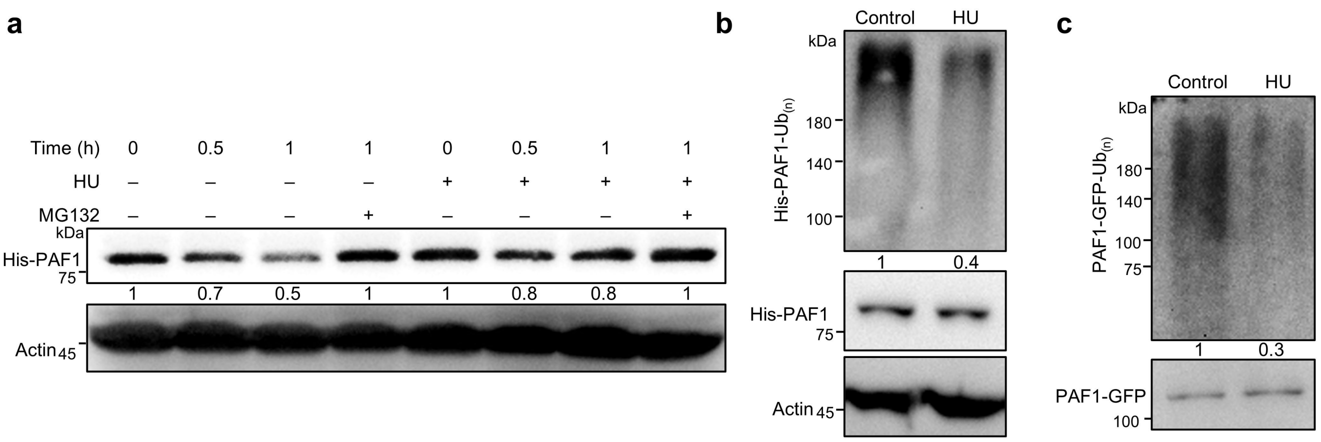


**Fig. S6 HU inhibits the polyubiquitination and degradation of PAF1.**

**a,** Semi-*in vitro* protein degradation assays. The recombinant His-PAF1 proteins were incubated with total protein extracts of Col-0 callus treated with or without 5 mM HU for 4 h. **b,** Semi-*in vitro* ubiquitination assays. The recombinant His-PAF1 proteins coupled with Ni-agarose beads were incubated with the total protein extracts of HU-treated Col-0 callus in ubiquitination buffer for 4 h. **c,** *In vivo* ubiquitination assays. *The 35S:PAF1-GFP/*Col-0 transgenic Arabidopsis were treated with 50 μM MG132 and/or 5 mM HU for 4 h. The proteins immunoprecipitated by anti-GFP beads were subjected to immunoblot analysis.

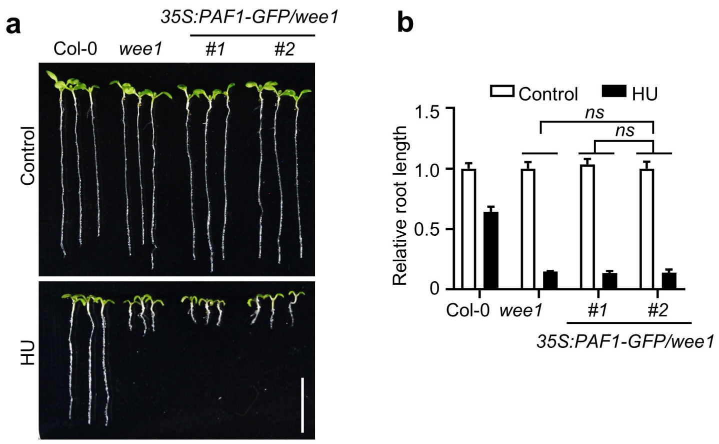


**Fig. S7 Overexpression of PAF1-GFP could not rescue the hypersensitivity of *wee1* to HU.**

**a,** Pictures of plants treated with HU. Plants were grown vertically on 1/2 MS medium with or without 1 mM HU for 7-8 days. Scale bar = 1 cm. **b,** The relative root length of the indicated plants. The data are represented as means ± SD (n = 10 plants) relative to the values obtained under the control conditions. The statistical significance was determined using two-way ANOVA analysis. *ns,* not significant.

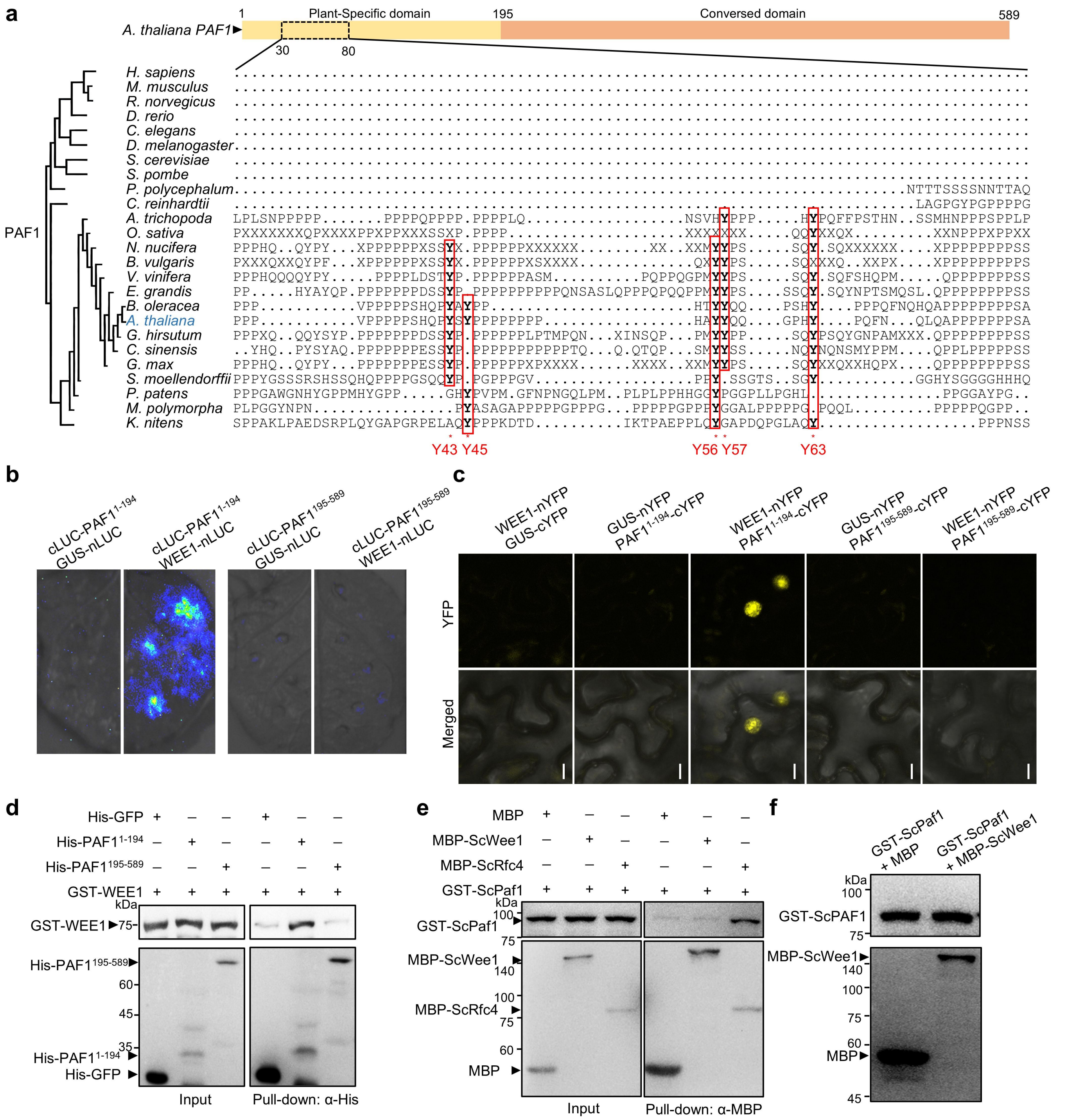

**Fig. S8 The regulatory mechanism of WEE1 on PAF1 is not conserved in yeast.**

**a,** Phylogenetic tree analysis and multiple sequence alignment on the amino acid sequences of PAF1 from 25 representative eukaryotes. The phosphorylation sites of PAF1 are highlighted in the red boxes. The data were obtained from PhyloGenes (https://phylogenes.arabidopsis.org/) with normal settings. **b,** Split luciferase assays. The proteins were fused to either the C- or N-terminal half of luciferase (cLUC or nLUC) and were transiently expressed in *N. benthamiana*. The luminescence detected by a charge-coupled device (CCD) camera indicates interaction. **c,** BiFC assays. The proteins were fused to either the C- or N-terminal half of YFP (cYFP or nYFP) and were transiently expressed in *N. benthamiana*. GUS serves as a negative control. The YFP fluorescence detected by confocal microscopy indicates interaction. Scale bars = 10 μm. **d and e,** *In vitro* pull-down assays. The GST or GST-WEE1 coupled with the glutathione beads was incubated with His-PAF1$^{1-194}$ or His-PAF1$^{195-589}$ (d). The MBP-ScWee1, MBP-ScRfc4, or MBP coupled with Dextrin beads was incubated with GST-ScPaf1 (e). **f,** *In vitro* protein phosphorylation assays. GST-ScPaf1 and MBP-ScWee1 or MBP were co-expressed in *E. coli*. The recombinant GST-ScPaf1 protein was purified with glutathione beads and analyzed by immunoblot analysis.

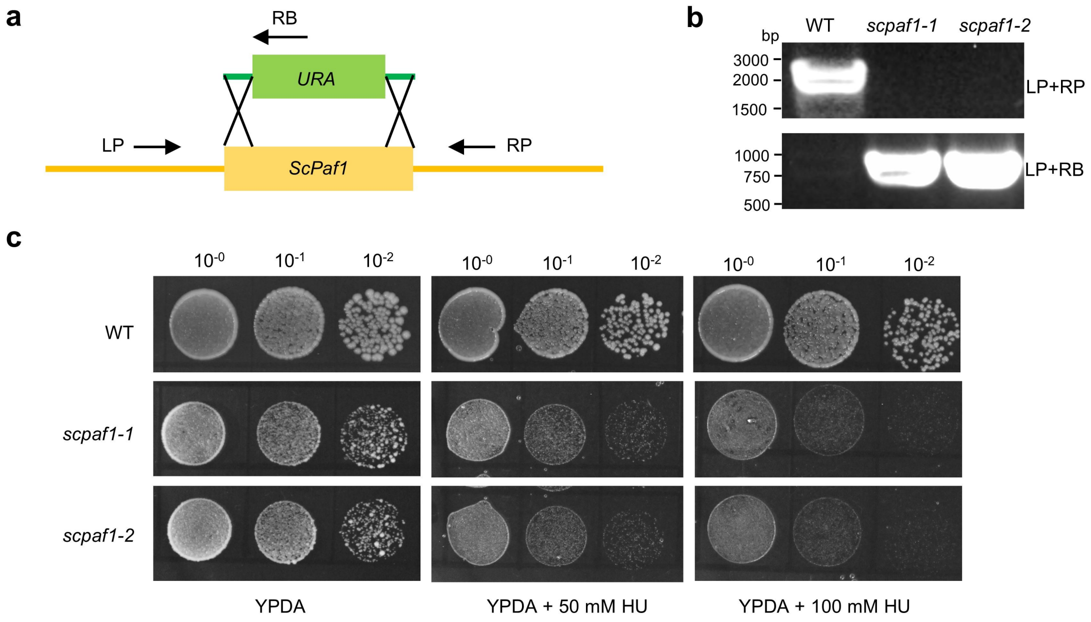


**Fig. S9 Yeast *scpaf1* mutant is hypersensitive to HU.**

**a,** Schematic representation of yeast *scPaf1* gene knockout. Homology between the flanking sequences at both sides of the *URA* gene and the *ScPaf1* gene allows the integration of the *URA* gene into the genome by the homologous recombination capability of yeast. **b,** Genotyping results of *scpaf1-1* and *scpaf1-2*. LP, RP, and RB are the primers shown in (a). **c,** Pictures of yeasts treated with HU. The WT, *scpaf1-1*, and *scpaf1-2* were grown for 4 days on YPDA medium with or without HU.

Proteins that copurified with PAF1-GS identified by mass spectrometry (Partial data )

| Protein | MS/MS count | Times detected |
|---|---|---|
| RFC1 | 772.3 | 2 |
| RFC2 | 491.4 | 2 |
| RFC4 | 320.8 | 2 |
| RFC3 | 202.9 | 3 |
| RAD17 | 96.1 | 2 |

**Fig. S10 The published PAF1 co-purification dataset (Antosz et al. 2017) shows that RFCs are co-purified with PAF1.**

Several subunits of the RFC complex were detected by triple-repeat mass spectrometry.

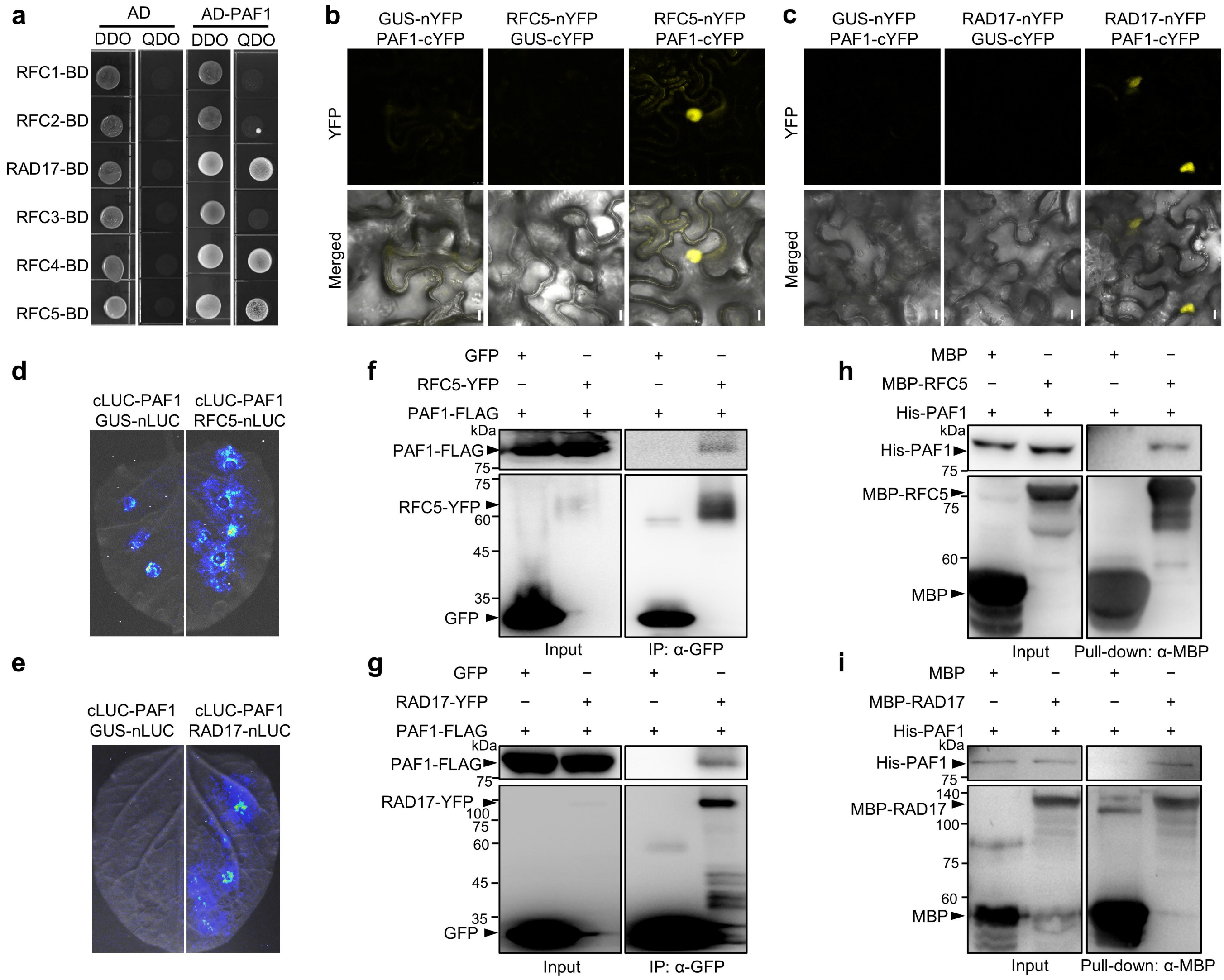


**Fig. S11 PAF1 interacts with RFC4, RFC5, and RAD17.**

**a,** Y2H assays. AD, activation domain. BD, DNA-binding domain. DDO, double dropout (SD/-Trp/-Leu) medium. QDO, quadruple dropout (SD/-Trp/-Leu/-His/-Ade) medium. The yeasts were grown for 3 days. **b and c,** BiFC assays. The proteins were fused to either the C- or N-terminal half of YFP (cYFP or nYFP) and were transiently expressed in *N. benthamiana*. GUS serves as a negative control. The YFP fluorescence detected by confocal microscopy indicates interaction. Scale bars = 10 μm. **d and e,** Split luciferase assays. The proteins were fused to either the C- or N-terminal half of luciferase (cLUC or nLUC) and were transiently expressed in *N. benthamiana*. The luminescence detected by a charge-coupled device (CCD) camera indicates interaction. **f and g,** Co-IP assays. PAF1-FLAG was co-expressed with GFP, RFC5-YFP, or RAD17-YFP in protoplasts. The immunoprecipitation was carried out using anti-GFP beads. **h and i,** *In vitro* pull-down assays. The MBP-RFC5, MBP-RAD17, and MBP coupled with Dextrin beads were incubated with His-PAF1.

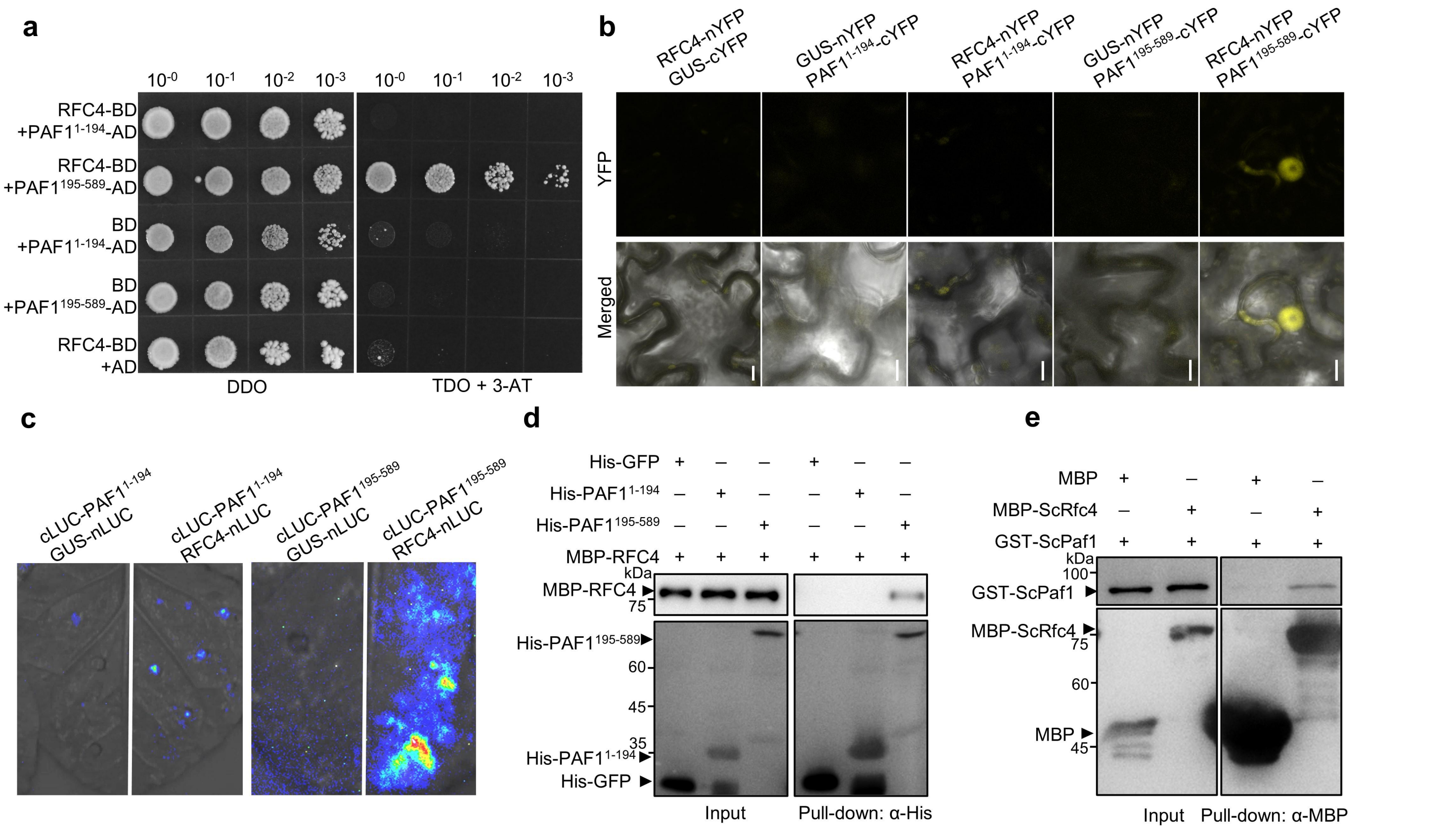


**Fig. S12 RFC4 interacts with the conserved domain of PAF1.**

**a,** Y2H assays. AD, activation domain. BD, DNA-binding domain. DDO, double dropout (SD/-Trp/-Leu) medium. QDO, quadruple dropout (SD/-Trp/-Leu/-His/-Ade) medium. The yeasts were grown for 3 days. **b,** BiFC assays. The proteins were fused to either the C- or N-terminal half of YFP (cYFP or nYFP) and were transiently expressed in *N. benthamiana*. GUS serves as a negative control. The YFP fluorescence detected by confocal microscopy indicates interaction. Scale bars = 10 μm. **c,** Split luciferase assays. The proteins were fused to either the C- or N-terminal half of luciferase (cLUC or nLUC) and were transiently expressed in *N. benthamiana*. The luminescence detected by a charge-coupled device (CCD) camera indicates interaction. **d and e,** *In vitro* pull-down assays. The His-PAF1$^{1-194}$, His-PAF1$^{195-589}$, or His-GFP coupled with the Ni-agarose beads were incubated with MBP-RFC4 (d). The MBP-ScRfc4 or MBP coupled with Dextrin beads were incubated with GST-ScPaf1 (e).

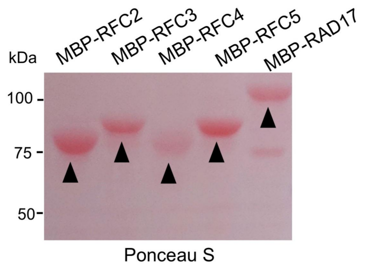


**Fig. S13 The purified MBP-RFC2/3/4 and MBP-RAD17 proteins.**

The purified MBP-RFC2/3/4 and MBP-RAD17 protein were used for the forked DNA-binding assays.
