## Supplementary material for "Plant-specific and conserved mechanisms of the polymerase-associated factor 1 complex in replication stress responses": Supplemantal information: Dataset S2.pdf

### Dataset S2. Multi-species sequence alignment analysis of PAF1

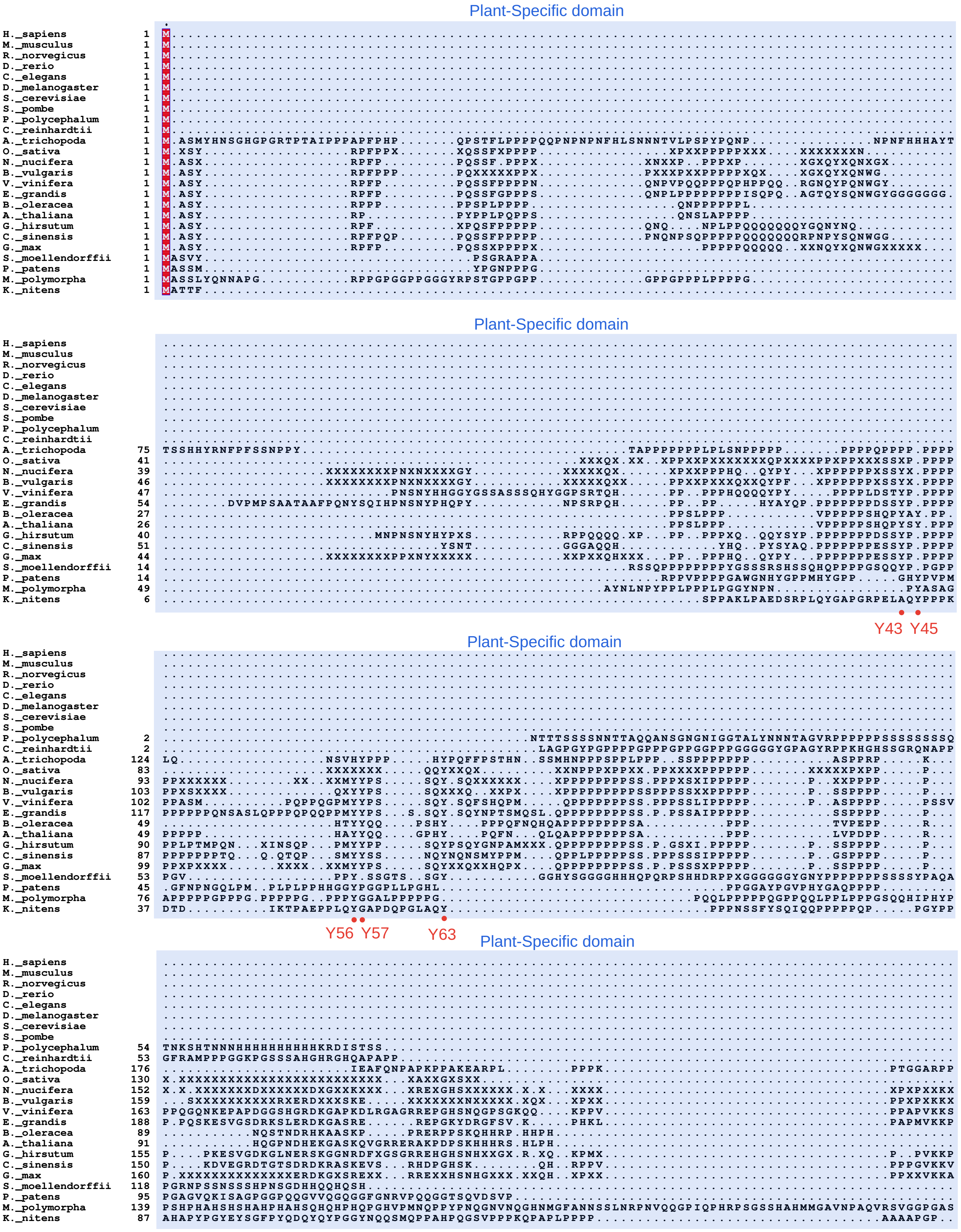

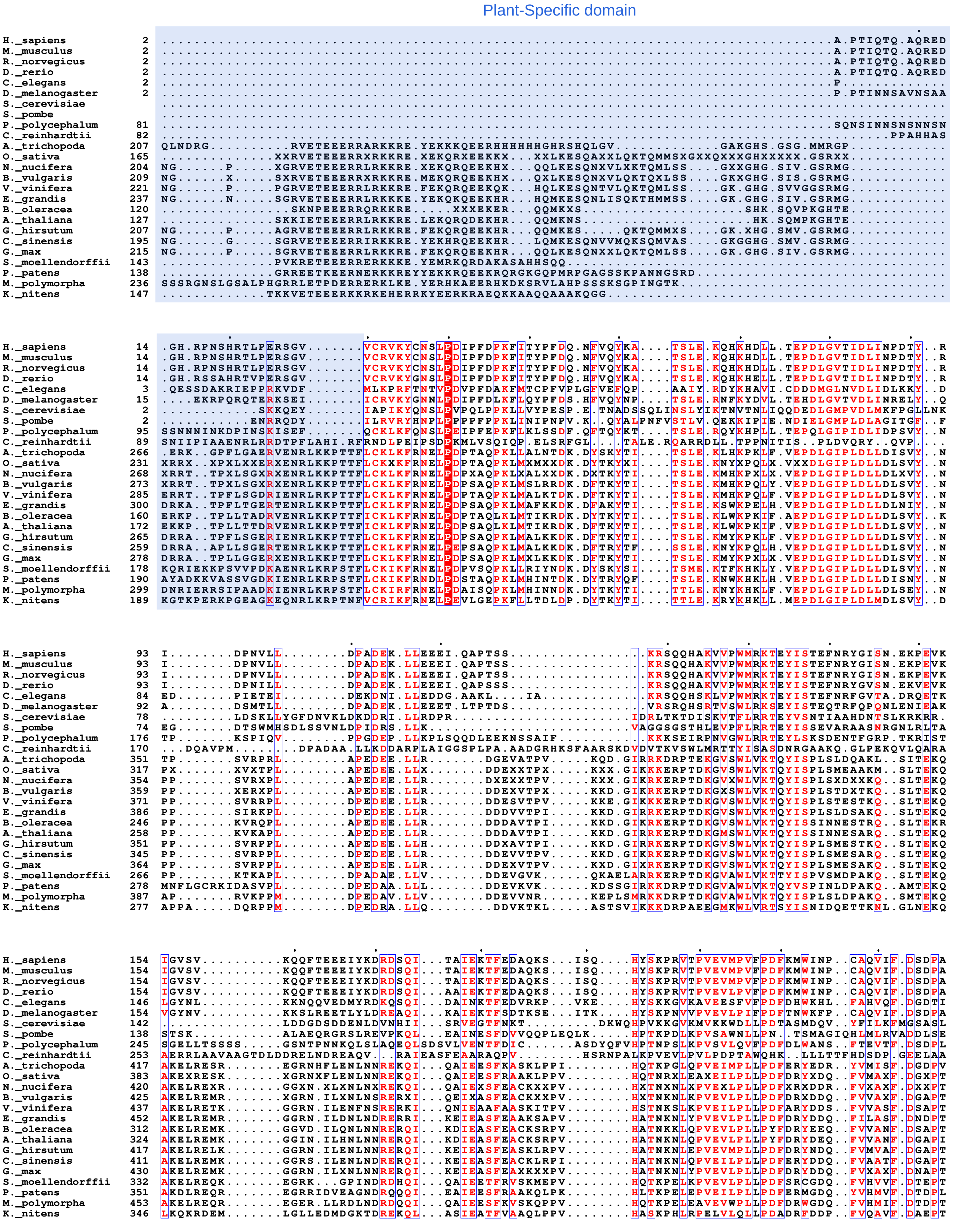
Plant-Specific domain

```
H._sapiens          228  PKDTSGAAALEMMSQAMIRGMMDEEGNQ....................FVAYFLPVEETL............KKRKRDQEE..EMDYAPDD.......
M._musculus         228  PKDTSGAAALEMMSQAMIRGMMDEEGNQ....................FVAYFLPVEETL............KKRKRDQEE..EMDYAPDD.......
R._norvegicus       228  PKDTSGAAALEMMSQAMIRGMMDEEGNQ....................FVAYFLPVEETL............KKRKRDQEE..EMDYAPDD.......
D._rerio            228  PKDVSAPAGVDMMSQAMIRGMMDEEGNQ....................FVAYFLPNEDTM............RKRKRDVEE..ELDYMPEE.......
C._elegans          220  TTEFEEEDERQQARESSVIKAMEFEDQK....................FAAVFVPTIGCL............TSFMDDLEL..ERPFDEDM.......
D._melanogaster     228  PAGKNVPAQLEEMSQAMIRGVMDESGEQ....................FVAYFLPTEQTL............EKRRTDFIN..GELYKEEE.......
S._cerevisiae       210  DTKEKKSLNT..................GIFRPVELEEDE........WISMYATDHKDS........AILENELEKGMDE..MDDDSHEGK......
S._pombe            214  RSHSYSSLVNLQEGHNLTKRHEV.....ALFMPSSAEGEE........FLSYYLPSEETA............EEIQAKVN...DASADVHE.......
P._polycephalum     328  DHFLPKDLRSDKVQEYASR..HNEVRNKAIVKGITDK...........FVYFITPDLKEN.......................YDVNNNND.......
C._reinhardtii      336  VVGQEAMAGLPSAKRPQLLVGHYLLKGFTHHISKAGQDRELKIMALLVREDVDRMVARTAAQVAQQEQEAAAAAAAEGREPAQAQAEPVLQPEDLEGD
A._trichopoda       493  AD........LEAYNKLDRATHDELESRAIMKSF.V.SGTDSSKPE.KFLAYLVPGPEEL............TK..........DMYDEHE.......
O._sativa           459  AD........SEXYXKLDXSXRDECESRAIMKXFSX.XGSDPXXXE.KFLAYMAPXPXEL............SK..........DIYXEXE.......
N._nucifera         495  AD........SEXYSKLDXSXRDXXESQAIMKSFXA.XGSDXXXPE.KFLAYMVPSPDEL............SK..........DXYDENE.......
B._vulgaris         500  AD........SEXYSKMDKSVRDXXESQAIMKSFVA.XGSDPXXPE.KFLAYMVPSXDEL............SK..........DIYDENE.......
V._vinifera         512  AD........SEIYSKLDKTVRDSHESQAILKSYMA.TGSDPSKPE.KFLAYMAPSPDEL............SK..........DIYDENE.......
E._grandis          527  AD........SEVYSKLEQSARDEYEAKAIMKSYEA.IGSDPSKPE.KFLAYMVPSADEI............SK..........DIYDVDE.......
B._oleracea         387  AD........SEFFGKLDSSIRDEHESRAVLKSYVI.AGSDNSNPE.KILAYMVPSLDEL............SK..........DMHDENE.......
A._thaliana         399  AD........SEFFGKLDPSIRDAHESRAILKSYVV.AGSDTANPE.KFLAYMVPSLDEL............SK..........DIHDENE.......
G._hirsutum         492  AD........SEIFSKLDXSVRDEHESRAIMKSYXA.ASSDPANPE.KFLAYMVPSLDEL............SK..........DMYDEHE.......
C._sinensis         486  AD........SEIYSKMDKSVRDAHESRAIMKSYVA.TGSDSANPE.KFLAYMVPSVNEL............SK..........DMYDENE.......
G._max              505  AD........SEXYXKLDXSVRDAYESRAXMKSYVA.TGSDPANPE.KFLAYMVPSPXEL............SK..........DXYDENE.......
S._moellendorffii   405  GD........SELYSKLDRNTRDYLESRAVVKSYSL..PKEDGKTE.KFLGYMVPKLDEA..........................HDDDD.......
P._patens           427  TD........APGLTDLDQDARLEIESRAIVKSFTI.ATSDESKPE.KFIGYMVPKPDEL............DR..........NFYDDEE.......
M._polymorpha       528  GE........SAQLSKLDIEMRDEIEARAIVKSYVLPPPSEDQKPE.KFVAYLIPKAEEL............DK..........DIYEEDD.......
K._nitens           422  AE........SERHRHYGADVREEMAARAMMKGFTI.PGDEGQAPE.KFIAYMVPSLEEV........................GADPEDR.......
```

```
H._sapiens          285  ...VYDYKIAREYNWN....VKNKASKGYEENYFFI.......FREGDGVYYNELETRVRLS.............KRRAKA..GVQSGTNA.......
M._musculus         285  ...VYDYKIAREYNWN....VKNKASKGYEENYFFI.......FREGDGVYYNELETRVRLS.............KRRAKA..GVQSGTNA.......
R._norvegicus       285  ...VYDYKIAREYNWN....VKNKASKGYEENYFFI.......FREGDGVYYNELETRVRLS.............KRRAKA..GVQSGTNA.......
D._rerio            285  ...VYEYKIAREYNWN....VKNKASKGYEENYFFI.......FRDADGVYYNELETRVRLS.............KRRAKV..GAQSSTNA.......
C._elegans          277  ...KYEFLLSREYTFK....MEHLPPRD...RDVFIM......YHRNNVFQYNEVDCNVKMT.............RKPKM.....ALSRKS.......
D._melanogaster     285  ...EYEYKIAREYNWN....VKTKASKGYEENYFFV.......MRQ.DGIYYNELETRVRLN.............KRRVKV...GQQPNNT.......
S._cerevisiae       266  ...IYKFKRIRDYDMKQVA........EKPMTELAIRL.....NDKDGIAYYKPLRSKIELR.............RRRVNDIIKPLVKEHD.......
S._pombe            277  ...PFVYNHFRNF.......DASMHVNSTGLEDLCLTFHTDKDHPEANQVLYTPIYARSTLK..............RRHVRAPVSLDAVDGIELSLR
P._polycephalum     383  ....FEID.DSQYKMQKVL.TSDIITDATSQNYFFLV........KNDAVYYNPLKNRVNL...............KKIKSKDDKLLVRTK.......
C._reinhardtii      433  YQWTKDYNYELKR..........DTVHYAL......RLDKDAARFYRMQGKLELRSWRDEEKLARDRSRKRVMGSANEEPAEEE.TDGSDQVQRPS
A._trichopoda       551  ...DIDFSWIREYHWD.VR.GDDAE....DPTTYLVN......FE.EGGARYLPLPTKLVLR.............RRR..I.DGRS...GH.D.IET
O._sativa           518  ...DXXYSWXREYHWD.VR.GDDXD....DPTTYLVX......FX.EXXAKYLPLPTKLVLQ.............KKR..AKEGRS...GD.E.XE.
N._nucifera         554  ...DXSYSWVREYHWD.VR.GDDAD....DPTTYLVX......FD.EXAARYLPLPTKLVLR.............KKR..AKEGRS...SD.E.VE.
B._vulgaris         559  ...DISYSWVREYHWD.VR.GEDAD....DPTTYLVS......FD.EXEARYLPLPTKLXLR.............KKR..AXEGRS...XD.E.VE.
V._vinifera         571  ...DTSYSWVREYHWD.VR.GDDAD....DPTTYLVS......FN.KTDARYLPLPTKLLLR.............KKR..AKEGRS...SD.E.VE.
E._grandis          586  ...EIGYSWLREYHWDFVRGGNDTE....EPSTYLVS......FD.EGEARYVPLPRKLNLR.............KKR..A.EGRS...SD.E.VE.
B._oleracea         446  ...DISYTWVREYLWD.IP.PKEPN....ERDTYLVS......FE.DGTASYLPLPTRLNMR.............KRR..AREGRA...SD.E.IE.
A._thaliana         458  ...EISYTWVREYLWD.VQ..PNAN....DPGTYLVS......FD.NGTASYLPLPMRLNLR.............KKR..AREGRS...SD.E.IE.
G._hirsutum         551  ...DISYSWVREYXWD.VR.GDDAN....DPTTYLVS......FD.EGEARYVPLPTKLNLR.............KKR..AREGRS...GD.E.IE.
C._sinensis         545  ...DVSFSWVREYHWD.VR.GDDAD....DPTTYLVS......FD.DDEARYVPLPTKLNLR.............KKR..AIEGRS...ND.E.VE.
G._max              564  ...DISYSWVREYHWD.VR.GDDAX....DPTTFLVS......FD.EXEARYLPLPTKLNLR.............KKR..AXEGRS...GD.E.VE.
S._moellendorffii   459  ...EIPYTWIREYQWD.VR.GEDAR......NNYVFC......FG.KDAVQYLPISSKLILH.............KKK..AREGRS...KD.D.VDN
P._patens           486  ...EIQYTWVREYHWD.IR.PDDSK.......NFVFC......FS.EDAVHYLPLGTKLVLQ.............KKK..AKDGKSKYDTE....DR
M._polymorpha       588  ...DIQYTWVREYHWD.LR.ADHQE......STYVFC......FG.EDSVTYLPLPSKLILQ.............KKK..AKEGRP...KE.EDFDN
K._nitens           479  ...EVEFDWVREYNWQLQP.TESGR......STFVLN......FE.DDQVAYLPIKNGLKLK.............KRK..ARESRQAPGSD.GEDDG
```

```
H._sapiens          347  ......LLVVK...........HRD.MNEKELEAQEARKAQLENHEPEEE..EEEEMETEEKEAGGSD.EEQE.......................KG
M._musculus         347  ......LLVVK...........HRD.MNEKELEAQEARKAQLENHEPEEE..EEEEMEAEEKEAGGSD.EEQE.......................KG
R._norvegicus       347  ......LLVVK...........HRD.MNEKELEAQEARKAQLENHEPEEE..EEEEMEAEEKEAGGSD.EEHE.......................KG
D._rerio            347  ......VLVCK...........HRD.MNEKELEAQEARKAQLENHEPED...EEEELDL.EKDMQEDSGEERE.......................KP
C._elegans          334  ......KLTLT...........YRN.PSELEQKDMNKREAELYEQPKTRKQEILEKIQEKKEE.................................GG
D._melanogaster     345  ......KLVVK...........HRP.LDSMEHRMQRYRERQLEVPGEEEEIVE.EVREEEQMQIIGETEKTSEDAAVGAQAASGADSPAQVARDRQS
S._cerevisiae       328  ....IDQLNVT...........LRN.PSTKEANIRDKLRMKFDPINFATVDE..............................................
S._pombe            350  DLNDEESLQLKRARYDTFGLGNIKD.LEEEEEKLRSVEGSLNEELSEEEKPAESREQL........................................
P._polycephalum     445  .IGRPESITYK...........KRP.QNSKELDAKEIIINDLLQEEKAPFSN..............................................
C._reinhardtii      512  KIHVT...........TRD.LNDRELEDRESKRKELMPL...........................................................
A._trichopoda       611  HYPVPSRVTVR...........RRS............TVATNELKES..GRAPS......................................MHDHSI
O._sativa           578  XFPVPSRITVR...........KXX............XVXXXELMEX..XXSSX......................................XHXGXX
N._nucifera         614  HYPIPSRVTVR...........CRP............EVAAIELQEX..XXYXX......................................XXKGXX
B._vulgaris         619  HFPVPXKXTVR...........RRX............XVSXIEXKET..ESYXX......................................SXXGXX
V._vinifera         631  HFPVPSKVTVR...........QRP............NVAAIELKDE..EVYSS......................................SKRGV.
E._grandis          647  HFARPAGVTVR...........RRT............IPATIELKEP..GVYSN......................................S.KRSA
B._oleracea         506  HLPVPSRVTVR...........NRS............SVSIIEHKDS..MVNTN......................................S.RNGA
A._thaliana         517  HFPVPSRVTVR...........RRS............TVSVIEHKDS..GVYSS........................................RVGA
G._hirsutum         611  HFPIPARITVR...........RRS............TVAAIELKEP..EVYXX......................................S.RGGM
C._sinensis         605  HFPIPSSIAVR...........RRA............NVTAIELKEQ..GAYSN......................................S.KGNS
G._max              624  HFPAPSRVTVR...........RRX............XVAAIELKDS..GXYSX......................................S.RGNX
S._moellendorffii   518  QFLVPSSVTVR...........RRDLTEEEEDQRESMRRIFMDGRLTETAIEEEAP..........................................
P._patens           545  DFPVPASVTVI...........RRDPTQDEEDIREDAREKFMEGGDT...................................................
M._polymorpha       648  QFPVPSRVTVH...........RRDLTDEELRARDEAKHAFMEGRARGAPDTGR............................................
K._nitens           543  AFRQPSRVTVQ...........KRS.QEEIEEDIRVAKIARIEEEERLEAEREER...........................................
```

```
H._sapiens          401  .....SSSEKEGS...EDEHSG.....SESEREEGDRDEASDKSGSGED...................ESSEDEA.RAARDKEEIFGSDADSEDDAD
M._musculus         401  .....SSSEKEGS...EDEHSG.....SESDREEGDRDEASDKSGSGED...................ESSEDEA.RAARDKEEIFGSDADSEDDAD
R._norvegicus       401  .....SSSEKEGS...EDERSG.....SESDREEGDRDEASDKSGSGED...................ESSEDEA.RAARDKEEIFGSDADSEDDAD
D._rerio            400  .....SDSEN.................SESES...EREEEERPADEDEEEEEDEESVKRRRERKSSGSESGDD...RQARDEEEIFGSDDDSEEEEE
C._elegans          381  .....DSSDQSSDSDDDKPQKSR......SDSSSDVSSDDDSPRKKEPTVDSD............................................
D._melanogaster     423  .....RSRSRTRSGSSSGSGSG.....SGSRASSRSKSGSRSGSGSRSRTNSPAGSQKS.....GSRSRSVSRSRSRSK......SGSRSRSRSRSK
S._cerevisiae       364  .....EDDEDEEQPEDVKKESEGDSKTEGSEQEGENEKDEEIKQEKENEQDEENKQDENRAADTPETSDAVHTEQKPEEEKETLQ............
S._pombe            407  .....ESAEQTNGVKPET.QAQNMSASESQANSPAPPVEEGNTQPSPVEQLQNE...........................................
P._polycephalum     484  .....KKLKRSNKLSDDE...............................................................................
C._reinhardtii           .................................................................................................
A._trichopoda       646  SKVAIPSKRGRSPVEDVHDDRRKVSRL...QD....MDDDQFSG..GEDM...............................................
O._sativa           613  SX...XXXRXXSXMDDXLXXXXXQRX...XD....X..DXXSG..EDDX...............................................
N._nucifera         649  SX...SXXXXXXTXEDGLVRXXKXXXX...QD....M..DHSSGA.EDDM...............................................
B._vulgaris         654  XX...SRXXXXXEXEDXXXXXXXXXXD...DD....X..DXXSGA.EDXM...............................................
V._vinifera         665  .....SSSKRGVDMEDGLGRSYKGVQD...QH....M..DQSSGA.EDEM...............................................
E._grandis          681  ST...SNLGALEDDEDDLGSPERVT.....QD..EHM..ERSSGA.EDDM...............................................
B._oleracea         540  SS...SKMRRL.EDEEGLGRSWKHGSE...QD....A..NQYSDN.NDDY...............................................
A._thaliana         550  SS...SKMRRL.EDEGGLGRSWKHEPE...QD....A..NQYSDGNEDDY...............................................
G._hirsutum         645  SS...SKXXRL.DAEDGLGRSHKLARH...QD....V..DQYSGA.EDDL...............................................
C._sinensis         639  SS...SKMGRV.DSQEDLERSHNGSRQ...QD....P..YQSSGA.EDDM...............................................
G._max              658  SX...SKMGXX.DXDDGLGRXHXXAXX...QD....X..XQSSGA.EDEM...............................................
S._moellendorffii   563  .....TTKSNKRPMEI..ERHSTKRRRPSYQE.......ETREASEDEDV...............................................
P._patens           581  .....YRRKRSRSPEDDIPVSTKRRGR............QDDYSDEEDEM...............................................
M._polymorpha       691  .....KRRRSPEAMMSSLKRKGVREEAVLNRE...........DVSDDEM...............................................
K._nitens           586  .....ERQEEEARAARAAERAARRDSDDEDSPKRGNESGEEDEAGQRDED...............................................
```

```
H._sapiens          465  SDDEDR...GQAQG.GSDNDSDSGSNG.GGQRSRSH....SRSASPFPSGSEHSAQEDGSEAA...ASDSSEA..DSDSD
M._musculus         465  SDDEDR...GQAHR.GSDNDSDSGSDG.GGQRSRSQSRSRSRSASPFPSGSEHSAQEDGSEAA...ASDSSEA..DSDSD
R._norvegicus       465  SDDEDR...GQAHR.GSDNDSDSGSDG.GGQRSRSQSRSRSRSASPFPSGSEHSAQEDGSEAA...ASDSSEA..DSDSD
D._rerio            469  EEEEGG...........................................ARRRSNSSSVQHSASERASDSSDA...SDSD
C._elegans          423  ..............................................................................SD
D._melanogaster     499  SGSRSRSGSRSGSGSRSPSRSRSGSPSG......................................SGSSSGSAS...DE
S._cerevisiae       444  ..............................................................................EE
S._pombe            455  ..............................................................................ED
P._polycephalum     497  ..............................................................................DI
C._reinhardtii      539  ........................................................................AE......
A._trichopoda       687  ..............................................................................SD
O._sativa           649  ..............................................................................SD
N._nucifera         686  ..............................................................................SD
B._vulgaris         691  ..............................................................................SD
V._vinifera         700  ..............................................................................SD
E._grandis          718  ..............................................................................SE
B._oleracea         576  ..............................................................................SE
A._thaliana         587  ..............................................................................SE
G._hirsutum         681  ..............................................................................SE
C._sinensis         675  ..............................................................................YD
G._max              694  ..............................................................................SE
S._moellendorffii   599  ..............................................................................SI
P._patens           614  ..............................................................................SV
M._polymorpha       725  ..............................................................................SV
K._nitens           631  ..............................................................................ED
```
